# Disentangling leaf structural and material properties in relation to their anatomical and chemical compositional traits in oaks (*Quercus* L.)

**DOI:** 10.1101/2022.10.21.513225

**Authors:** David Alonso-Forn, Domingo Sancho-Knapik, María Dolores Fariñas, Miquel Nadal, Rubén Martín-Sánchez, Juan Pedro Ferrio, Víctor Resco de Dios, José Javier Peguero-Pina, Yusuke Onoda, Jeannine Cavender-Bares, Tomás Gómez Álvarez Arenas, Eustaquio Gil-Pelegrín

## Abstract

The existence of sclerophyllous plants has been considered an adaptive strategy against different environmental stresses. As it literally means “hard-leaved”, it is essential to quantify the leaf mechanical properties to understand sclerophylly. However, the relative importance of each leaf trait on mechanical properties is not yet well established. The genus *Quercus* is an excellent system to shed light on this since it minimizes phylogenetic variation while having a wide variation in sclerophylly. Thus, leaf anatomical traits and cell wall composition were measured, analyzing their relationship with LMA and leaf mechanical properties in a set of 25 oak species. Outer wall contributed strongly to leaf mechanical strength. Moreover, cellulose plays a critical role in increasing leaf strength and toughness. The PCA plot based on leaf trait values clearly separated *Quercus* species into two groups corresponding to evergreen and deciduous species. Sclerophyllous *Quercus* species are tougher and stronger due to their thicker epidermis outer wall and/or higher cellulose concentration. Furthermore, section *Ilex* species share common traits regardless of they occupy quite different climates. In addition, evergreen species living in Mediterranean-type climates share common leaf traits irrespective of their different phylogenetic origin.

## 1 Introduction

Sclerophyllous leaves have been considered a universal adaptive strategy in response to different environmental stresses (Alonso-Forn *et al*. 2020). Thus, stress factors such as water deficit (e.g., Schimper, 1903; Oertli *et al*., 1990), nutrient shortage (Loveless, 1961; 1962; Beadle, 1966), low temperatures (Koppel & Heinsoo, 1994; Lamontagne *et al*., 1998) and physical damage (Chabot & Hicks, 1982; Grubb, 1986) would have a synergistic effect that may explain the variation in sclerophylly. This trait is a characteristic recognized by botanists according to a physiognomic criterion. Thus, botanists have described sclerophyllous leaves as hard, tough, stiff and leathery (Schimper, 1903; Seddon, 1974; Turner, 1994). Given that the concept of sclerophylly is a perception, it is difficult to obtain an accurate measure or objective classification of sclerophylly (Edwards *et al*., 2000). Moreover, as sclerophylly literally means "hard-leaved" it is essential to quantify the leaf mechanical properties to understand this trait.

Most ecophysiological studies use the leaf mass per unit area (LMA) as a quantitative proxy value for sclerophylly (e.g., Niinemets, 2001; Wright *et al*., 2004). LMA is a combination of leaf thickness (LT) and leaf density (LD), which in turn depend on a great variety of anatomical and compositional leaf traits such as vein density, vein diameter, number of cell layers, cell size, air space fraction or fiber content (Mediavilla *et al*., 2008; John *et al*., 2017; Sancho-Knapik *et al*., 2021). Sclerophylly has been well studied in the genus *Quercus*, and in this regard, Sancho-Knapik *et al*. (2021) recently found that LT influences the increase in LMA more than LD. While LMA provides a good proxy for leaf “hardness”, it is not necessarily a measure of leaf mechanical resistance (Onoda *et al*., 2011).

Direct approaches to quantify leaf sclerophylly require the measurement of leaf mechanical properties (Aranwela *et al*., 1999; Read & Sanson 2003; Onoda *et al*., 2011). These include the fracture-related properties, such as strength and toughness, which refer to the ability to resist an applied force and applied work, respectively (Cherrett, 1968; Williams, 1954; Coley, 1983; Choong *et al*., 1992). A higher structural strength (i.e., maximum force to fracture), a higher structural toughness (i.e., work to fracture) and their thickness-normalized properties namely specific strength and specific toughness have been previously associated with a higher leaf sclerophylly (Edwards *et al*., 2000; Wright & Cannon, 2001; Read & Sanson 2003). Leaf mechanical properties can be analyzed in relation to their underlying components, which is analogous to the decomposition of LMA into LD and LT (Lucas *et al*., 2000; Kitajima & Poorter, 2010; Lusk *et al*., 2010). In the punch-and-die test, structural mechanical properties such as maximum force or work to fracture per unit fracture length (punch strength and punch toughness) can be decomposed into material properties (specific punch strength and specific punch toughness) and LT (Onoda *et al*., 2008; Onoda *et al*., 2011; Westbrook *et al*., 2011). Material properties can be further analyzed in relation to tissue density or cell wall fiber content (percentage of hemicellulose, cellulose, and lignin per unit leaf dry mass or per unit volume).

The relative importance of each element of leaf tissue (e.g., cuticle, epidermis, palisade mesophyll, spongy mesophyll and vascular bundle extension) on the mechanical properties is not well established because their relative volume fraction have different influences on LMA and mechanical properties. Actually, several studies found that some anatomical variables strongly influenced leaf toughness through the reinforcement of certain structures with little effect on the amount of accumulated biomass per unit surface area (Edwards *et al*., 2000; Westbrook *et al*., 2011; Onoda *et al*., 2012; 2015). Onoda *et al*. (2015) showed that some structural traits such as the cuticle thickness have a significant impact on mechanical strength and toughness. The main objectives of the present study were to determine the leaf anatomical and compositional traits underlying a higher leaf strength and leaf toughness. The objective was achieved by measuring leaf anatomical traits and cell wall fiber composition, analyzing their relationship with both LMA and leaf mechanical properties.

The genus *Quercus* is an excellent system to perform this study, as it not only minimizes phylogenetic variation (compared to studies performed across diverse species), but also displays a wide variation in leaf sclerophylly across species (Gil-Pelegrín *et al*., 2017). In particular, section *Ilex* species has been considered as pre-adapted to dry climates (He *et al*., 2014). Although their ancestors occupied tropical and subtropical wet forests in the Himalaya-Hengduan mountains, some species exhibit xeromorphic-like traits (Het *et al*., 2014; Jiang *et al*., 2019). These traits would have allowed them to cope with hot-dry seasonal conditions that occur in the Mediterranean-type climates (Martín-Sánchez et al. 2022).

The present study was performed on a set of 25 oak species (*Quercus* spp.) with different leaf habit (deciduous and evergreen) grown in a common garden in Northeastern Spain. We analysed 41 leaf structural, morphological, physicochemical and anatomical traits on these species (for species names see **Fig. 5)**. Specifically, we address the following hypotheses. (i) While structural mechanical properties are better correlated with leaf anatomical traits, leaf material properties are better correlated with cell-wall composition. (ii) Sclerophyllous leaves are stronger and tougher due to their thicker epidermis outer wall and/or higher cellulose concentration.

## 2 Material and Methods

### 2.1 Plant Material

For this study, leaves were sampled from a living collection of 25 oak species, maintained in the experimental fields from CITA de Aragón (41°390N, 0°520W, 200 m a.s.l., Zaragoza, Spain). This common garden features Mediterranean climatic conditions with a mean annual temperature of 15.4 °C and total annual precipitation of 298 mm. Oak trees were *ca*. 20 years old; they were drop irrigated twice per week and pruned if it was necessary. Current year, fully developed, mature leaves were collected from south-exposed branches of one individual per species during the early morning. Leaves were stored in sealed plastic bags and carried to the laboratory in portable coolers. One set of 10 leaves was used for punch and die tests to measure the mechanical properties, leaf strength and toughness. A second set of nine leaves was aimed to phenolic compounds analysis. A third set of 10 leaves was utilized to obtain LMA. A fourth set of 10 leaves was used for leaf fiber content analysis, and a fifth set of 10 leaves was used to obtain the morphological and anatomical traits.

### 2.2 Mechanical properties: punch and die test

Punch and die tests consisted of punching a hole through the leaf lamina. As the punch contacts the leaf surface, the tip applies pressure until it overcomes the tensile strength of the leaf, causing a fracture. As a result, a leaf hole is produced and the compressive forces on the punch are released. A flat-ended and sharp-edged cylindrical punch made of steel of 2 mm diameter with a clearance of 0.05 mm was built and mounted onto the moving head of the mechanical tester Mach-1 V500C MA001 system (Biomomentum, Inc., Québec, Canada). Similar to Read and Sanson (2003) a die designed to fit the punch was located in the threaded base of the machine. A typical trial allowed the punch to penetrate the die to a depth of 2 mm at a speed of 30 mm min^−1^. Data for all punch tests were collected at a rate of 100 Hz and were used to generate force-displacement curves (Fig. **S1a**). Before every set of measurements, a blank test was performed as a calibration in order to account for measuring the load forces due to the background friction caused by the proximity of the die walls to the punch tip. Leaf thickness (LT) was also measured with a micrometer (GT-H10L, Keyence, Osaka, Japan) attached to an amplifier unit (GT-75AP, Keyence, Osaka, Japan) just before each punch and die test.

To minimize variation in results due to differences between leaves of the same species, the leaf tissue tested was standardized: major veins were avoided and all trials were made halfway between the secondary veins delimiting the upper and lower borders of the intercostal panel. When the size of the species under study was big enough, four tests were conducted in each of the five leaves selected. Otherwise, two tests were performed and ten leaves selected (Fig. **S1b**). All the mechanical tests were taken at room temperature with leaves full hydrated.

The analysis of the curves allowed to obtain the following parameters: maximum force (F_max_) defined as the highest load value, punch strength (PS) calculated as F_max_ divided by the circumference of the punch, and the leaf toughness or work of fracture (WF), calculated as the area below the curve between the initial contact of the punch with the leaf and F_max_ (Fig. **S1a**). The starting point of the curve was counted from 10% of F_max_, to avoid the effect of the leaf three-dimensional structure. Afterwards, the specific punch strength (SPS) and the specific work of fracture (SWF) were expressed per unit area of fracture (see all abbreviations and units in **Table 1**).

**Table 1.** List of mechanical and chemical properties, leaf morphological, and leaf anatomical traits with their abbreviations and units. List of leaf mechanical properties, anatomical and compositional traits with their abbreviations and units.

| Abbr. | Parameter | Units | Abbr. | Parameter | Units |
| --- | --- | --- | --- | --- | --- |
| BSE_cp | Bundle sheath extension cover percentage | % | LT | Leaf thickness | $\mu\text{m}$ |
| BSE_d | Bundle sheath extension density | $\text{mm mm}^{-2}$ | N.LD | Nitrogen per unit volume | $\text{mg cm}^{-3}$ |
| BSE_w | Bundle sheath extension width | $\mu\text{m}$ | Ncw_Ntotal | Nitrogen in cell wall:Nitrogen | |
| C.LD | Carbon per unit volume | $\text{mg cm}^{-3}$ | Ntotal | Nitrogen content | $\text{g } 100^{-1} \text{ g}^{-1}$ |
| CC | Cellulose content | $\text{mg g}^{-1}$ | PM_cl | Palisade mesophyll cell length | $\mu\text{m}$ |
| Ccw_Ctotal | Carbon in cell wall:Carbon | | PM_cw | Palisade mesophyll cell width | $\mu\text{m}$ |
| CD | Cellulose per unit volume | $\text{mg cm}^{-3}$ | PM_nl | Palisade mesophyll number of layers | |
| Ctotal | Carbon content | $\text{g } 100^{-1} \text{ g}^{-1}$ | PMT | Palisade mesophyll thickness | $\mu\text{m}$ |
| Ctotal_Ntotal | Ratio carbon:nitrogen | | PS | Punch strength | $\text{kN m}^{-1}$ |
| HC | Hemicellulose content | $\text{mg g}^{-1}$ | SM_p | Spongy mesophyll porosity | % |
| HCD | Hemicellulose per unit volume | $\text{mg cm}^{-3}$ | SMT | Spongy mesophyll thickness | $\mu\text{m}$ |
| LCC | Lignin and cutin content | $\text{mg g}^{-1}$ | SPS | Specific Punch Strength | $\text{MN m}^{-2}$ |
| LCD | Lignin and cutin per unit volume | $\text{mg cm}^{-3}$ | SWF | Specific Work to Fracture | $\text{kJ m}^{-2}$ |
| LD | Leaf density | $\text{mg cm}^{-3}$ | UE_latw | Upper epidermis lateral wall | $\mu\text{m}$ |
| LE_latw | Lower epidermis lateral wall | $\mu\text{m}$ | UE_lu | Upper epidermis lumen cell | $\mu\text{m}^2$ |
| LE_lu | Lower epidermis lumen cell | $\mu\text{m}^2$ | UE_lul | Upper epidermis lumen length | $\mu\text{m}$ |
| LE_lul | Lower epidermis lumen length | $\mu\text{m}$ | UE_luw | Upper epidermis lumen width | $\mu\text{m}$ |
| LE_luw | Lower epidermis lumen width | $\mu\text{m}$ | UE_ow | Upper epidermis outer wall | $\mu\text{m}$ |
| LE_ow | Lower epidermis outer wall | $\mu\text{m}$ | UET | Upper epidermis thickness | $\mu\text{m}$ |
| LET | Lower epidermis thickness | $\mu\text{m}$ | WF | Work of fracture | $\text{J m}^{-1}$ |
| LMA | Leaf mass per unit area | $\text{g m}^{-2}$ | | | |

### 2.3 Leaf mass per area, compositional content, lignification and arid intensity

To calculate LMA, one disc (12.6 mm^2^ in area) per leaf was obtained between the secondary veins of 10 leaves per species. Discs were oven-dried for 3 days at 70 °C and afterwards, they were weighed to obtain their dry mass. LMA was then calculated as the ratio between the dry mass and the disc area. Additionally, leaf density (LD) was calculated as the ratio between LMA and LT (Niinemets, 1999; Sancho-Knapik *et al*., 2021).

For fiber content calculation, 10 fresh leaves were oven-dried for 3 days at 70 °C. Then, the petiole and mid-rib were removed. The rest of the plant material was ground and values of hemicellulose content (HC), cellulose content (CC) and lignin + cutin content (LCC) were obtained by quantifying neutral detergent fiber (NDF) and acid detergent lignin (ADL) following the method of Goering and Van Soest (1970). The amount of total foliar nitrogen and carbon (Ntotal and Ctotal, respectively) in dry leaves was measured using an organic element analyzer (Flash EA 112, Thermo Fisher Scientific Inc., Waltham, MA, USA). The cellular composition was obtained after the neutral detergent fiber (NDF) procedure according to the method of Goering and Van Soest (1970). The nitrogen content of the cell wall fraction was further estimated using the elemental analyzer described above.

To detect the presence of lignified anatomical structures, 15-20 μm cross-sections were cut with a microtome (HM 350 S, MICROM GmbH, Walldorf, Germany). Then, cross-sections were stained with a drop of phloroglucinol-HCl solution or Wiesner stain, prepared as a mixture 2:1 of 3 % phloroglucinol in absolute ethanol and concentrated HCl (Pradhan & Loqué, 2014). Sections were observed under a light microscope (OPTIKA B-600TiFL, Optika Microscopes, Ponteranica, Italy) where lignified tissues appeared as fuchsia in color. A few drops of Safranin and AstraBlue 0.1% double stain were added to confirm tissue lignification with a second method. After 30 seconds, the samples were rinsed with distilled water and observed under light and epifluorescence microscopy (OPTIKA B-600TiFL, Optika Microscopes, Ponteranica, Italy) using green filter.

We selected a climatic variable to characterise aridity stress intensity: the arid intensity, defined as the sum of (2*t*_m_ – *p*_m_) for months with 2*t*_m_ > *p_m_* and *t*_m_ > 10°C. Being *t*_m_, the mean monthly temperature and *p*_m_, the mean monthly precipitation. Data was obtained from Sancho-Knapik *et al*. (2019).

### 2.4 Anatomical traits

Vein morphological parameters were determined in a set of five mature leaves per species following the method described in Scoffoni *et al*. (2011) with some modifications. Leaf sections obtained between secondary veins were chemically cleared with 5% NaOH in an aqueous solution, washed with a bleach solution, dehydrated in an ethanol dilution series (70, 90, 95 and 100 %) and stained with safranin. Afterwards, one image (40x) per sample was taken using a camera (Canon EOS M100) coupled to a microscopy (OPTIKA B-600TiFL, Optika Microscopes, Ponteranica, Italy) and venation-related traits were measured in three fields per leaf using the ImageJ software. Bundle sheath extension density (BSE_d) was calculated as the ratio between the sum of all bundle sheath extension lengths and sampled area. The cover percentage of the leaf surface occupied by bundle sheath extension (BSE_cp) and the mean width of the bundle sheath extension (BSE_w; Fig. **1c**) were obtained using the Image J software.

**Fig. 1.**
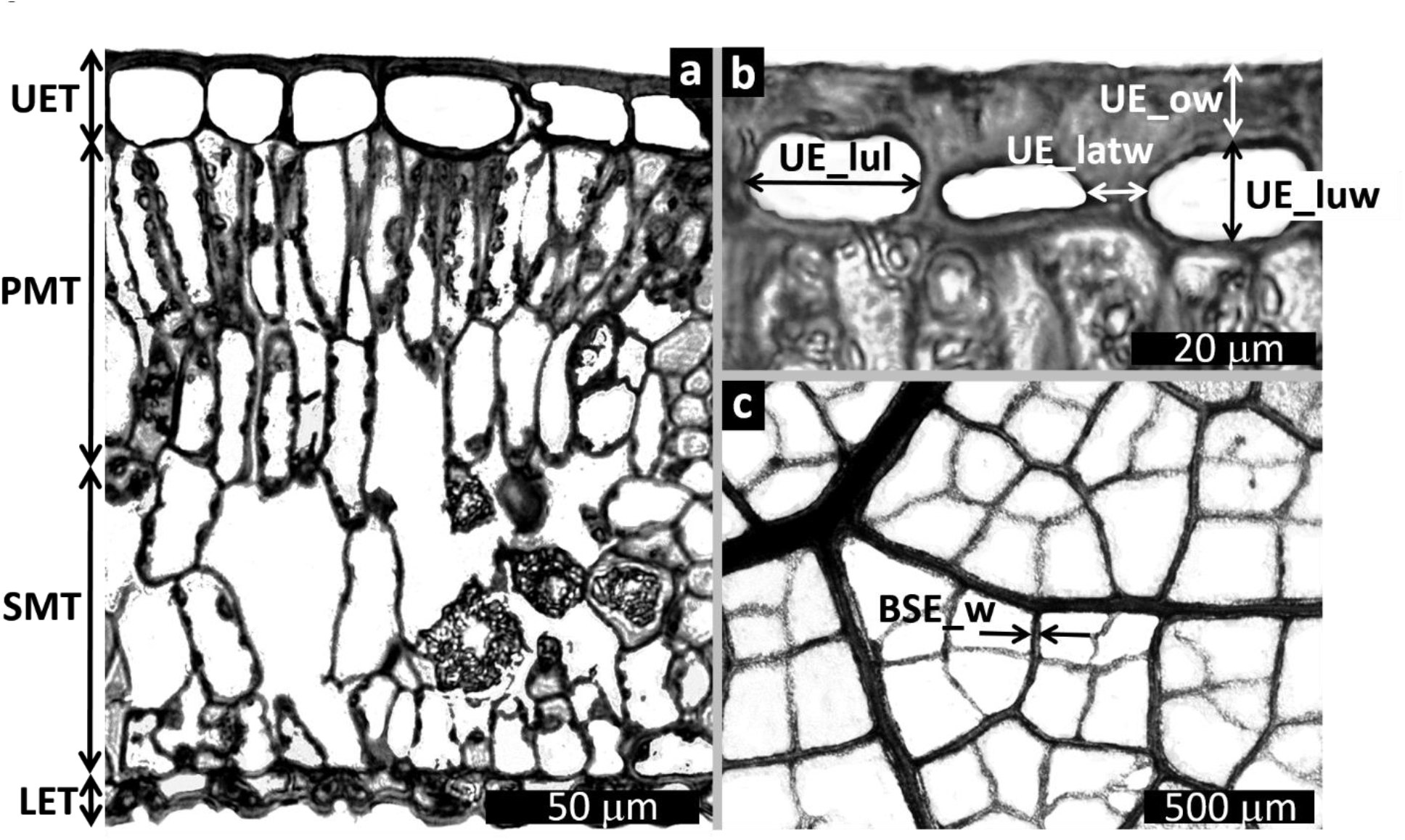
Leaf mesophyll cross section of *Quercus myrsinifolia* (a), detail of the upper epidermis of *Q. chrysolepis* (b), and detail of leaf venation of *Q. shumardii* (c), showing diverse anatomical leaf traits measured in this study. Trait notation as in **Table 1**.

Finally, anatomical sections of five leaves per species were obtained by gradual dehydration with ethanol, propylene oxide as a transition agent and inclusion in Araldite. One mesophyll image (200x) per leaf section was taken using a camera (Canon EOS M100) coupled to a microscope (OPTIKA B-600TiFL, Optika Microscopes, Ponteranica, Italy) and the following parameters were measured in five fields of each image: palisade and spongy mesophyll thickness (PMT and SMT, respectively; Fig. **1a**), number of cell layers in the palisade mesophyll (PM_nl), palisade mesophyll cell width and length (PM_cw and PM_cl, respectively), spongy mesophyll porosity (SM_p), upper and lower epidermis thickness (UET and LET, respectively; Fig. **1a**), upper and lower epidermis outer wall (UE_ow and LE_ow, respectively; Fig. **1b**), upper and lower epidermis lumen width and length (UE_luw, UE_lul, LE_luw and LE_lul, respectively; Fig. **1b**), upper and lower epidermis lateral wall (UE_latw and LE_latw, respectively; Fig. **1b**) and cell lumen size (UE_lu and LE_lu, respectively). See all parameters, abbreviations and units in **Table 1**.

### 2.5 Structural equation models (SEM)

The correlation analyses identified the independent variables most correlated to punch strength and specific punch strength (Fig. **S2**). With this information in mind, we proposed two mechanistic models including the most representative variables to estimate the network of correlations between traits related to leaf mechanical properties, which were assessed with structural equation models (SEM). In the first model, the leaf structural properties (i.e., leaf thickness, palisade mesophyll thickness, spongy mesophyll thickness, upper - lower epidermis thickness and upper - lower epidermis outer wall thickness) were related to punch strength (Fig. **3a**). In the second model, the leaf material properties (leaf density, cellulose content, hemicellulose content, and lignin and cutin content) were related to the specific punch strength (Fig. **3b**).

### 2.6 Leaf construction cost

Leaf samples from the seven studied species were oven dried at 70 °C for 3 d until constant mass, ground and homogenized. Total leaf N concentration was determined with an Organic Elemental Analyzer (Flash EA 112, Thermo Fisher Scientific Inc., MA, USA). Ash concentration was determined gravimetrically after combustion of duplicated samples for at least 4 h at 500 °C. Heat of combustion was determined in triplicate samples of 18-24 mg with an adiabatic bomb calorimeter (Phillipson Gentry Instruments, Inc., USA) with correction for ignition wire melting (Villar & Merino, 2001), following the procedure of Phillipson (1964). Leaf construction cost (CC) (g glucose g^−1^) was calculated according to Williams *et al.* (1987) as:

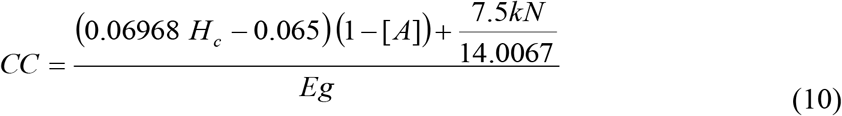

where *H*_c_ is the ash-free heat of combustion (kJ g^−1^), [*A*] is the ash concentration (g ash g^−1^ dry mass), *N* is the tissue nitrogen concentration (g g^−1^ dry mass), *Eg* is the growth efficiency (0.89 for woody leaves; Williams *et al.*, 1987) and *k* is the oxidation state of the nitrogen source (+5 for nitrate or −3 for ammonium). In well-aerated Mediterranean soils, nitrate is the main source of nitrogen and thus, *k* = +5. However, given that nitrate reduction in leaves can occur at the expense of reductive equivalents generated at light with no apparent cost, we note that the nitrogen contribution to the construction cost also depends on the share of nitrate reduction above- and belowground (Niinemets, 1997; Niinemets, 1999).

### 2.7 Statistical analysis

Data passed Shapiro–Wilk and Bartlett tests for normality and equality of variances, respectively. Interspecific differences in leaf traits were evaluated by one-way ANOVA. All analyses were performed in R (R Core Team, 2021). To summarize the multivariate relationships among anatomical traits and *Quercus* species, a principal components analysis (PCA) with two components was carried out. This PCA was conducted using the FactoMineR: PCA package (Josse & Husson, 2008). To find correlations between all studied parameters, a Pearson’s correlation matrix was performed using ‘corrplot’ package (Wei & Simko, 2021). SEM were implemented using ‘lavaan’ package (Rosseel, 2012). Standardized major axis (SMA) regressions (Warton *et al*. 2006) were fitted to summarise the “scaling” relationship between two variables. To ensure our results did not artificially arise from lack of phylogenetic independence, we additionally performed phylogenetic generalized least squares analysis assuming that trait evolution mimics Brownian motion and using the phylogeny from Hipp *et al*. (2020).

## 3 Results

### 3.1 Leaf mass per area and leaf traits behind mechanical properties

Leaf mass per area (LMA), which is a combination of leaf thickness (LT) and leaf density (LD), ranged from 64.2 to 223.9 g m^−2^. LT ranged from 169 to 487 μm and showed a good correlation (*r* = 0.84, *p* < 0.001) with LMA. However, LD showed less variation from 273 to 579 mg cm^−3^ and had a moderate correlation (*r* = 0.52, *p* = 0.008) with LMA. Oak species in this study showed a wide range in the values of the leaf mechanical properties measured. Punch strength (PS) and specific punch strength (SPS) ranged from 0.59 to 4.47 kN m^−1^ and from 2.63 to 11.11 MN m^−2^, respectively (Fig. **2**), work of fracture (WF) from 0.07 to 1.31 J m^−1^, specific work of fracture (SWF) from 0.33 to 3.26 kJ m^−2^. The species that showed the highest values of these mechanical parameters were *Quercus chrysolepis*, *Q. phillyreoides*, *Q. coccifera* and *Q. ilex* subsp. *rotundifolia*. In contrast, the species with the overall lowest values were *Q. lobata*, *Q. robur* and *Q. cerris*. In general, measured mechanical properties showed a good correlation (*r* > 0.7, *p* < 0.001) with LMA (Fig. **2**) even when phylogeny was considered (r > 0.73, p < 0.01; Fig. **S4**). PS showed a strong positive correlation (*r* = 0.75, *p* < 0.001) with LT, but a weak correlation (*r* = 0.4, *p* = 0.045) with LD. Moreover, PS can be mathematically expressed as the product of SPS, and LT with LT having a relative contribution of 0.51 to PS. SPS was better correlated (*r* = 0.62, *p* < 0.001) with LD than with LT (*r* = 0.38, *P* > 0.05; Fig. **2**). Furthermore, leaves showed higher PS and WF at a given LMA as the scaling coefficients of the standardized major axis slopes were significantly steeper than 1 (1.47 for PS and 2.06 for WF; see Fig. **S3**). As has been seen in LMA and leaf mechanical resistance measurements, a wide range of variation was found in the compositional and anatomical variables (see table **S1** and TRY dataset).

**Fig. 2.**
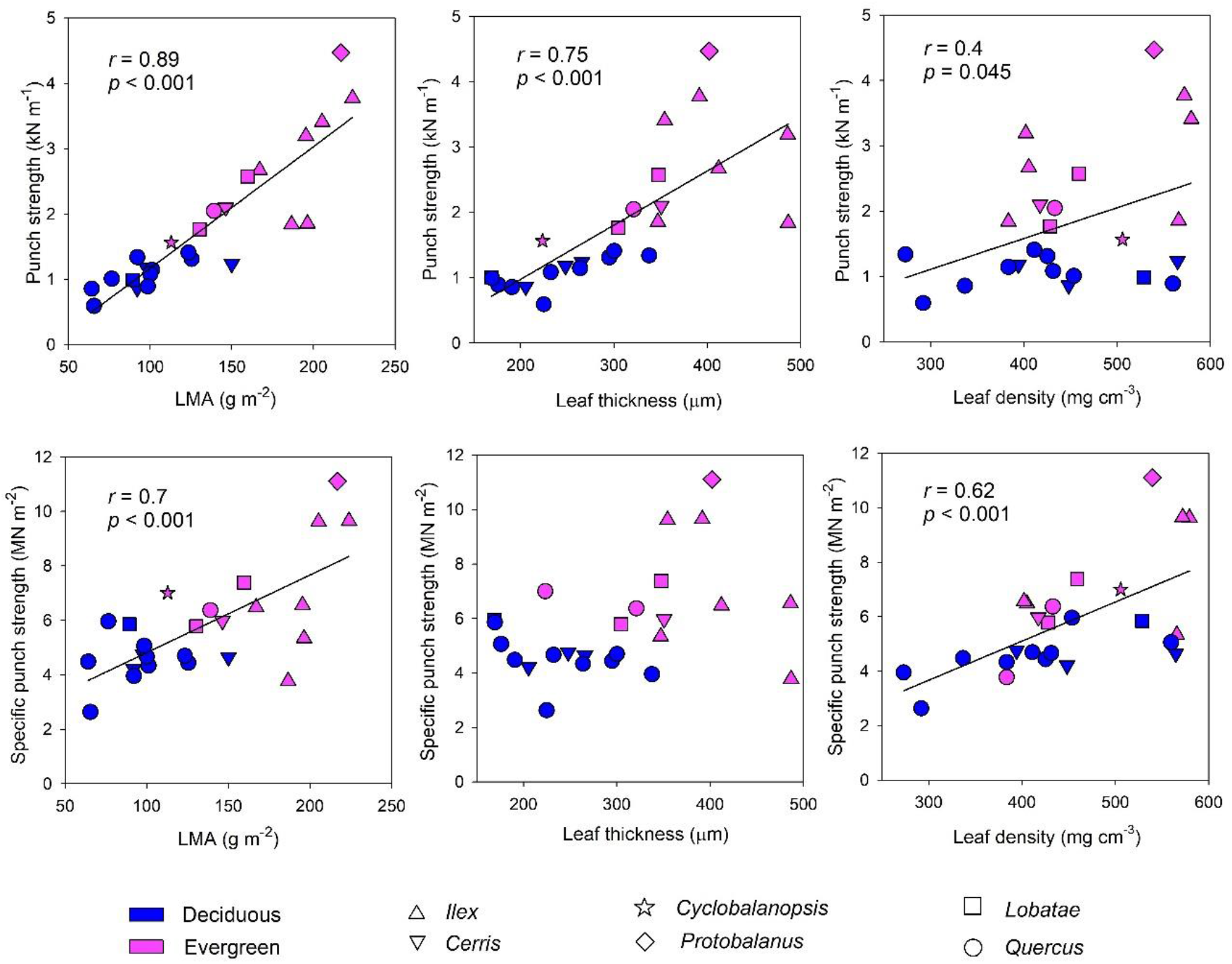
Main relationships between physical parameters (punch strength and specific punch strength) and leaf mass per area, leaf thickness, and leaf density, for deciduous (DEC; blue) and evergreen (EVE; pink) *Quercus* species. Each point belongs to a *Quercus* species and represents its mean value. Black continuous line is the correlation considering all species when correlation was significative (*p* < 0.05).

Both material and mechanical properties were positively correlated with cellulose concentration (CC; *r* > 0.72, *p* < 0.001), palisade and spongy mesophyll thicknesses (PMT and SMT, respectively; *r* = 0.50 and 0.86, repectively *p* < 0.05) including the components palisade mesophyll cell width and number of layers (PM_ct and PM_nl, respectively; *r* = 0.43 to 0.79, *p* < 0.05), upper epidermis and lower epidermis outer wall and lateral wall (UE_ow, UE_latw and LE_ow, LE_latw; *r* = 0.59 to 0.86, *p* < 0.01) and bundle sheet extension density, width and cover percentage (BSE_d, BSE_w and BSE_cp, respectively; *r* = 0.42 to 0.79, *p* < 0.05; Fig. **S2**). Upper epidermis lumen cell (UE_lul) were negatively correlated with mechanical properties (*r* < −0.61). The rest of the anatomical and compositional traits showed less significant correlation coefficients (*r* < 0.5) or did not show significant correlations with the mechanical properties (Fig. **S2**). The presence of lignified anatomical structures was detected in all species, specifically in the bundle sheath. However, clearly lignified epidermis only appeared in the most sclerophyllous species (eg. *Q. chrysolepis*, *Q. phyllireoides* and *Q. coccifera*).

Leaf carbon concentration and nitrogen concentration in cell wall (Ncw_Ctotal and Ncw_Ntotal) were not related to leaf mechanical traits. However, leaf carbon concentration and C/N ratio did have a moderate correlation with PS (C.LD and C_N; *r* = 0.40, *p* = 0.049 and *r* = 0.41, *p* = 0.044, respectively).

### 3.3 Structural equation models (SEM)

In the designed paths for SEMs, leaf traits were divided, on the one hand, into anatomical traits (thickness of the different tissue layers) related to LT and SPS (Fig. **3a**), and, on the other hand, into compositional traits (cell wall components) related to LD and SPS/LD (Fig. **3b**). In the anatomical traits model, both LT and SPS showed a high contribution to PS (*r* = 0.49 and *r* = 0.7, respectively). Regarding anatomical traits, LT was highly explained by PMT (*r* = 0.46) while SMT and ET showed a non-significant association with LT (Fig. **3a**). On the other hand, SPS was strongly associated with the upper epidermis outer wall thickness (UE_ow) (*r* = 0.78) while it did not influence LT. Furthermore, the sum of upper and lower epidermis thickness (ET) had a moderate influence on SPS (*r* = 0.33). UE_ow and ET together explained much of the variation in SPS (*R*^2^ = 0.54).

**Fig. 3.**
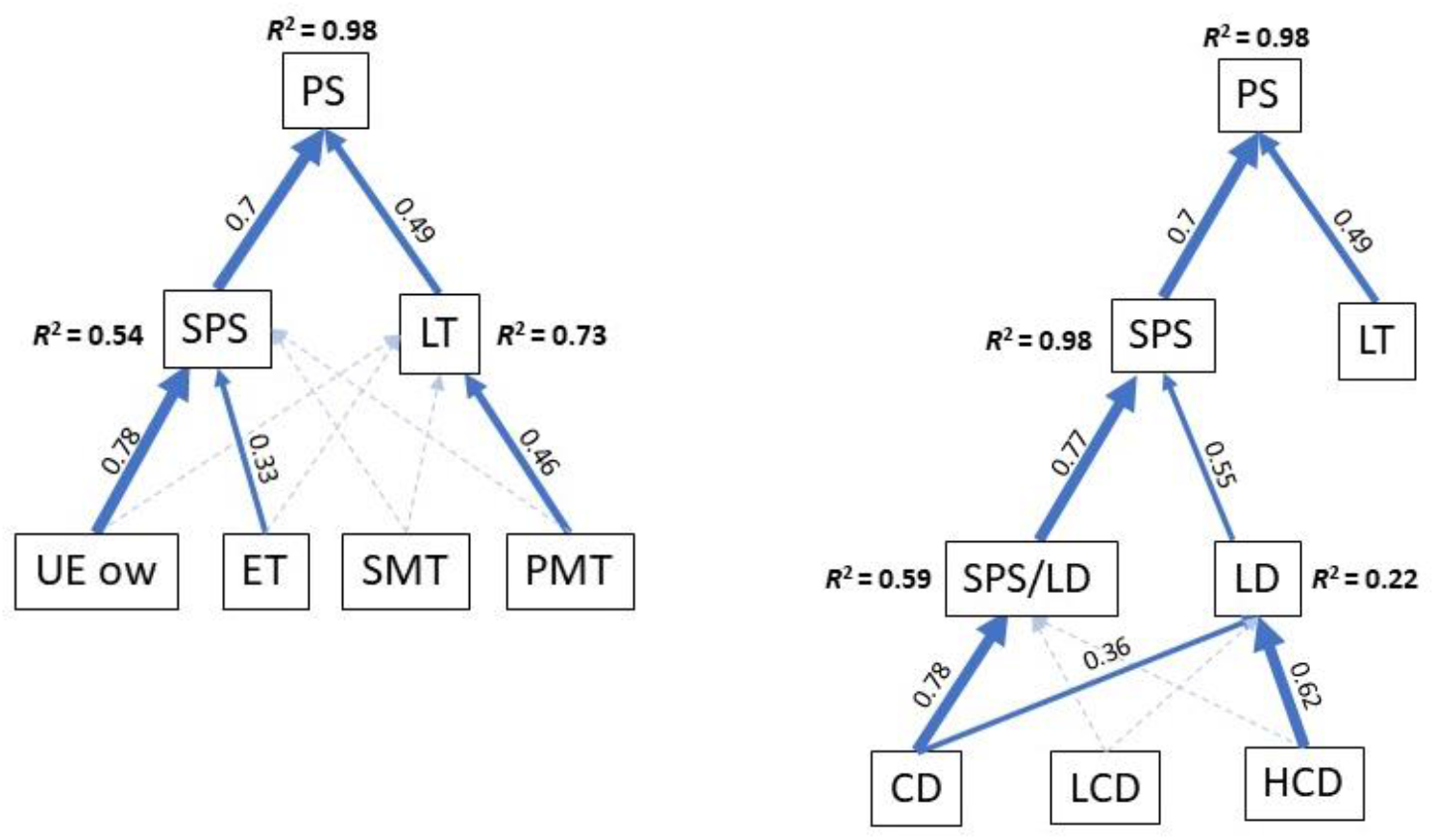
Structural equation models (SEM) for the contribution of (a) anatomical and (b) compositional variables to leaf strength. punch strength (PS) and b) specific punch strength (SPS). The figure shows the full model, with significant correlations (*p* < 0.05) highlighted by solid arrows. PS, punch strength; SPS, specific punch strength; LT, leaf thickness; LD, leaf density; PMT and SMT, palisade and spongy mesophyll thickness, respectively; ET, summatory of upper and lower epidermis thickness; UE_ow, upper epidermis outer wall; CD, cellulose content per unit volume; HCD, hemicellulose content per unit volume; LCD, lignin and cutin content per unit volume. Numbers denote standardized path coefficients.

In the compositional traits model, SPS was strongly associated with both LD (*r* = 0.55) and SPS/LD (*r* = 0.77). As expected, LD was explained by a combination of compositional traits, especially hemicellulose content per unit volume (HCD, *r* = 0.62) and cellulose content per unit volume (CD, *r* = 0.36). SPS/LD was explained by a strong (*r* = 0.78) direct influence of CD (Fig. **3b**).

### 3.4 Principal component analysis and phylogeny

A principal component analysis (PCA) based on a selection of the strongest correlations between variables in the correlation matrix (Fig. **S2**) and considering the SEM, was performed. This PCA showed that the first main component (explaining the 62.4% variability) grouped PS, LMA, CD, PMT, SMT, UE_ow and BSE_w, whereas the second main component (explaining the 15.7% variability), grouped LD and HCD (Fig. **4**). The scores of the studied *Quercus* species in the PCA biplot indicated that the mechanical, compositional and anatomical traits analyzed clear differentiated deciduous and evergreen species although could not differentiate the species according to the climate (Fig. **4**).

**Fig. 4.**
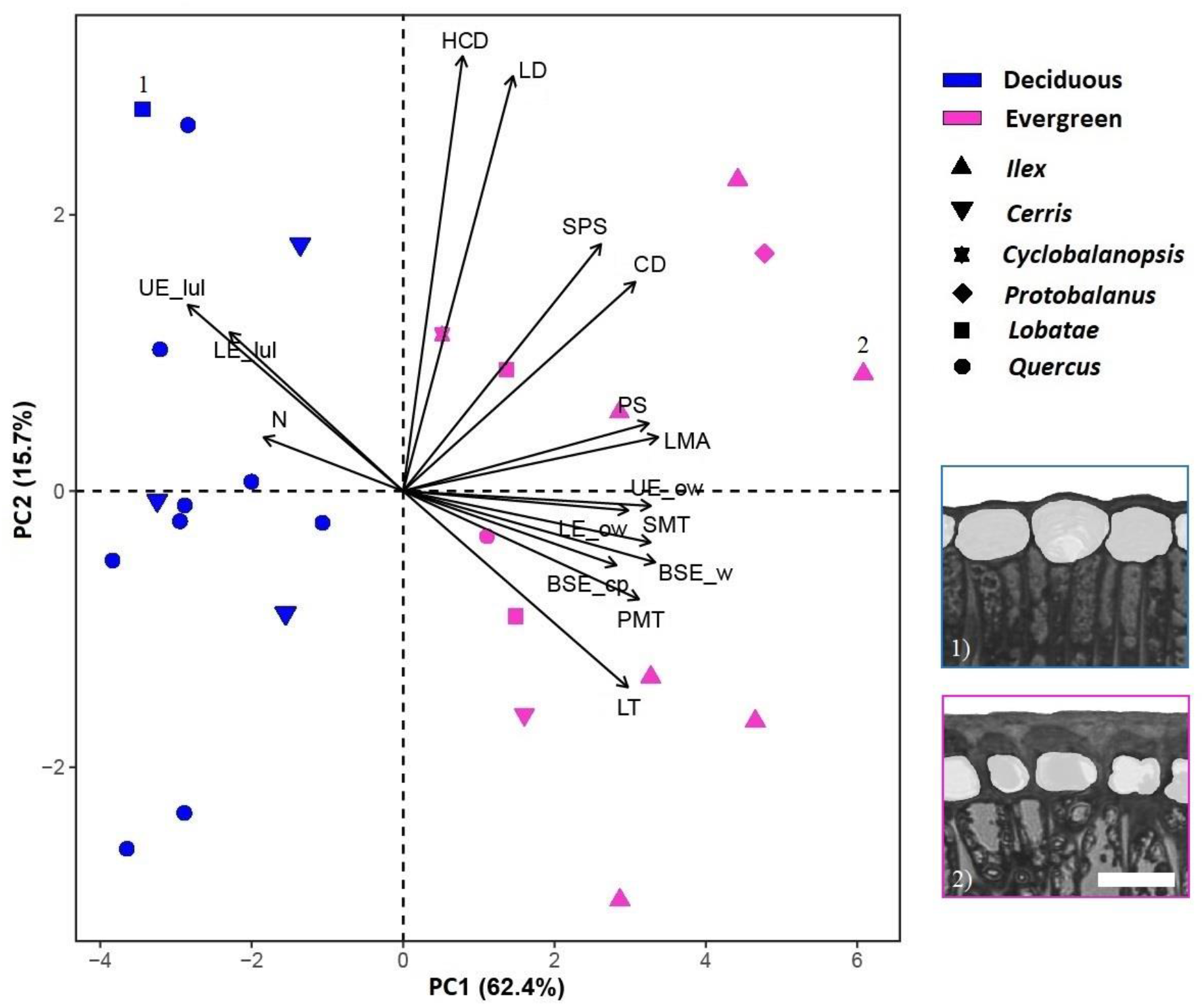
Principal component analysis (PCA) of leaf variables (black lines) in 25 *Quercus* species (dots). Each symbol corresponds to a genus *Quercus* section. Variables: PS, punch strength; SPS, specific punch strength; LMA, leaf dry mass per unit area; LT, leaf thickness; LD, leaf density; CD, cellulose content per unit volume; HCD, hemicellulose content per unit volume; LCD, lignin and cutin content per unit volume; PMT, palisade mesophyll thickness; SMT, spongy mesophyll thickness; UET, upper epidermis thickness; LET, lower epidermis thickness; UE_ow, upper epidermis outer wall; LE_ow, lower epidermis outer wall; BSE_cp, bundle sheath extension cover percentage; BSE_w, bundle sheath extension width; N, Nitrogen content; UE_lu, upper epidermis cell lumen size; LE_lu, lower epidermis cell lumen size. Images of the epidermis of (1) *Q. shumardii* and (2) *Q. phillyreoides*. Scale bar, 20 μm.

Regarding the phylogeny, all the species of the *Quercus* section, except *Q. mohriana*, which is evergreen, tend to have low LMA, PS, cellulose content, thin mesophylls and UE_ow (Fig. **5a**). In contrast, all species of the *Ilex* sect. had a strong coordination with those traits related to sclerophylly as high LMA, PS, cellulose content, thick mesophyll and UE_ow, although they inhabit very different climates. *Q. chrysolepis* (sect. *Protobalanus*), presented sclerophyllous traits. Among the studied sect. *Lobatae* species, those evergreens (*Q. agrifolia* and *Q. wislizeni*) had a strong coordination with sclerophyllous traits, while *Q. shumardi* had less relation to sclerophyllous traits and a strong coordination with LD and hemicellulose content. In the case of *Q. myrsinifolia* (sect. *Cyclobalanopsis*), it had moderate coordination with both main components. In the case of the sect. *Cerris*, the only species with sclerophyllous traits was *Q. suber*. Regarding climate, no significant relationship was found between arid intensity and the main components of PCA (Fig. **5b**; Fig. **S5**).

**Fig. 5.**
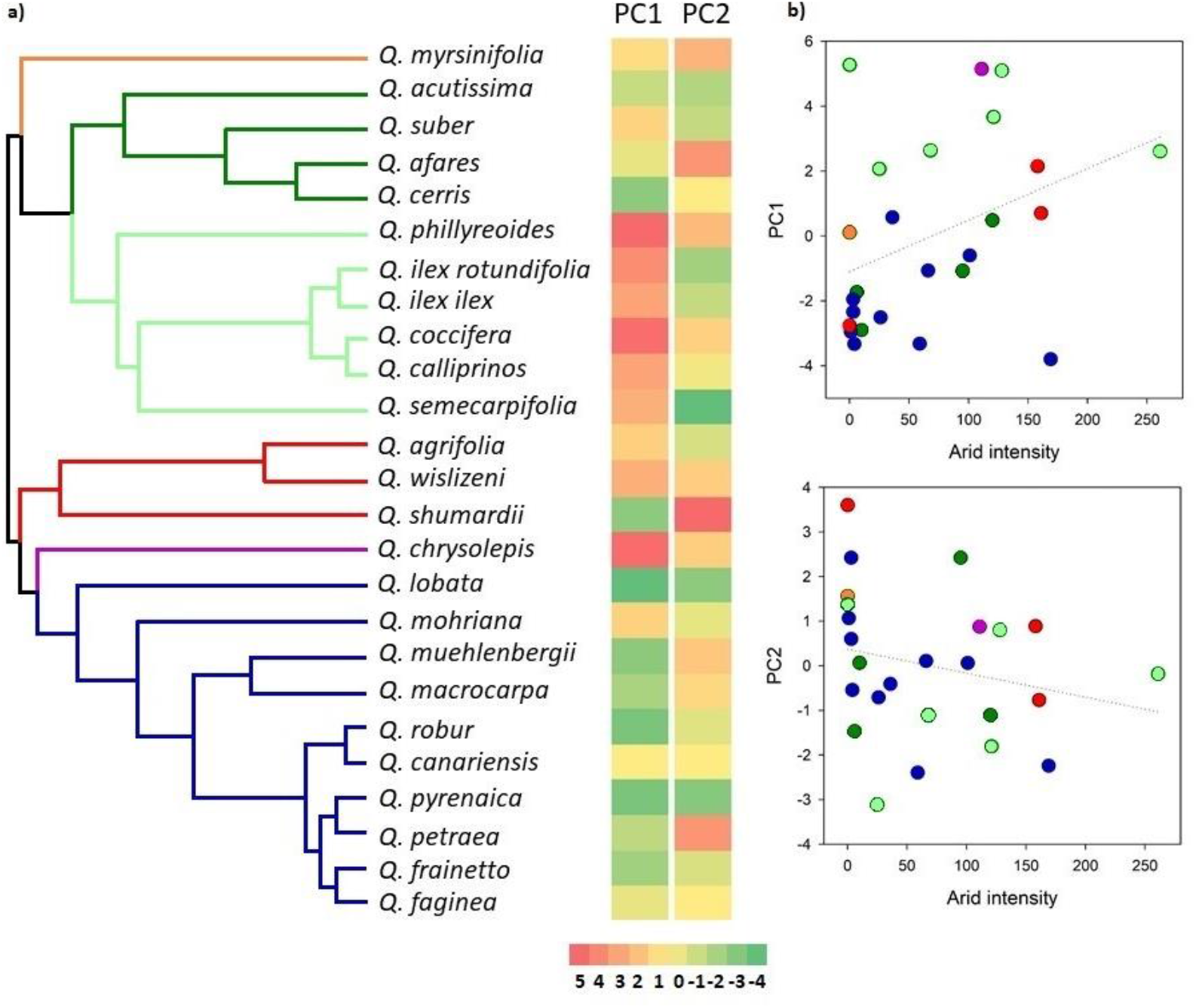
a) *Quercus* phylogenetic tree based on Hipp *et al*. (2020). Colors in the tree represent *Quercus* sections; in orange, section *Cyclobalanopsis*; dark green, section *Cerris*; light green, section *Ilex*; red, section *Lobatae*; fuchsia, section *Protobalanus*; blue, section *Quercus*. PC1 and PC2. b) Relationship between PC1 and PC2 and Arid intensity (AI). The colors of the circles refer to the phylogenetic tree sections. It should be noted that the lower the AI value, the higher the degree of aridity.

### Differences in construction cost and underlying traits among the evergreen oaks

Leaf construction cost of the seven species ranged from 1.43 ± 0.05 g glucose g^−1^ (*Q. ilex* subsp. *rotundifolia*) to 1.61 ± 0.06 g glucose g^−1^ (*Q. agrifolia* and *Q. coccifera*), while the mean leaf construction cost for the seven strudied species was 1.53 ± 0.07 g glucose g^−1^ (**Table S2**). All the studied species showed construction cost values in the range published for sclerophyllous evergreen leaves (Villar & Merino, 2001). The ash fraction does not contribute to the construction cost, while the nitrogen concentration increases the construction cost due to extra costs required for nitrogen assimilation. However, the energy content (*H*_c_) was the main determinant of the leaf construction cost; therefore, the highest *H*_c_ values corresponded to the highest construction cost values (**Table S2**). Thus, *Q. suber*, with the highest leaf N concentration, but also with the highest ash content, and low energy content had the lowest leaf construction cost (**Table S2**).

## 4 Discussion

In this study, we analysed 41 leaf mechanical, morphological, physicochemical and anatomical traits of 25 *Quercus* species grown in a common garden in Northeastern Spain to determine the leaf traits underlying sclerophylly. Although most ecophysiological studies have used leaf mass per unit area (LMA) as a proxy value for sclerophylly, sclerophylly actually means “hard-leaved,” implying that mechanical properties of leaves are important in understanding this attribute (Alonso-Forn *et al*., 2020). As previously observed (Wright *et al*., 2004; Onoda *et al*., 2011), our results showed that, mechanical properties were strongly correlated with LMA in studied *Quercus* species.

The inability of LMA to account adequately for the wide variation in punch strength in our dataset reveals that there are variations in the mechanical properties that do not contribute to an increase in the accumulated mass per surface area. In this sense, we found a variation range in punch strength (PS) measures of 7.5-fold and in work to fracture (WF) of 18-fold, while LMA only varied 3.5-fold. Moreover, leaves showed higher PS and WF at a given LMA as the scaling coefficients of the standardized major axis slopes were significantly steeper than 1 (1.47 for PS and 2.08 for WF; see Fig. **S3**). Although the increase in LMA in *Quercus* has previously been attributed mainly to an increase in leaf thickness (LT) (Sancho-Knapik *et al*., 2021), we found that when accounting for the effect of LT, the specific work to fracture (SWF) still increased with LMA (Fig. **2**; **S3**). These results suggest that LMA could have more implications besides leaf strength. Although high LMA species can have lower photosynthetic rate per unit leaf area (Wright et al. 2004), Sancho-Knapik *et al*. (2021) suggested that accumulation of photosynthetic tissues enable plants to generate higher assimilation rates when facing certain environmental limitations. Moreover, some sclerophyllous evergreen oaks, despite their larger LMA, were able to achieve area-based net CO2 assimilation values equivalent to congeneric deciduous species due to an increased chloroplast surface area exposed to intercellular air space (Peguero-Pina *et al*., 2017; Onoda *et al*., 2017).

According to our models, the upper epidermis outer wall had a strong and direct contribution to the leaf mechanical strength (measured as specific punch strength, SPS) (Fig. **3a**) which is consistent with previous studies (Onoda *et al*., 2012; Westbrook *et al*., 2011; He *et al*., 2019). Thus, an increase in the thickness of this layer appears to lead to a tougher epidermis without great changes in the accumulated mass per unit leaf surface area. Onoda *et al*. (2012) showed that leaves with relatively thicker cuticles had higher leaf mechanical resistance because the cuticle is made of very stiff material. Although a thicker cuticle may imply a higher investment (Poorter & Villar, 1997), it does not result in substantial changes in leaf thickness, total accumulated mass per surface nor construction cost (Villar & Merino, 2001; **Table S2**). In this sense, although *Q. chrysolepis* a notably higher (>60%) LMA than *Q. agrifolia*, their construction costs are very similar (**Table S2**). In addition, its extra cost can be offset by its greater mechanical resistance to biotic and abiotic stresses which might confer longer leaf lifespans among evergreen species (Krauss *et al*., 1997; Riederer & Schreiber, 2001; Wright *et al*., 2004; Carver & Gurr, 2006; Onoda *et al*., 2012).

The cellulose concentration per unit volume had a strong direct contribution to SPS/LD (Fig. **3b**) which indicates that cellulose plays a critical role in increasing leaf strength and toughness, a result also found in tropical plants (Kitajima & Poorter, 2010). In the correlation analysis (Fig. **S2**), cellulose concentration was tightly correlated with the bundle sheath extension width and cover percentage, and the outer wall thickness, suggesting that these anatomical components, which are rich in cellulose, play key roles in mechanical resistance.

The PCA plots based on foliar trait values indicated a clear separation of two groups along the first component axis corresponding to evergreen and deciduous species (Fig. **4**). Foliar mechanical properties, including punch strength, LMA, cellulose content, mesophylls thickness upper epidermis outer wall thickness, Bundle Sheath Extension width and cover percentage separated these contrasting leaf habits. Deciduous oaks from *Quercus*, *Cerris* and *Lobatae* sections, showed common leaf attributes regardless of their phylogenetic lineage. This group of species is characterized by softer and thinner leaves, narrower bundle sheath extension, less coverage of bundle sheath extension, lower cellulose content, as well as thinner cuticles and outer walls that are not lignified. Thus, *Q. robur* (section *Quercus*) from central Europe and *Q. shumardii* (section *Lobatae*) from eastern North America share these leaf traits (Fig. **5**). Moreover, these leaf features do not seem to depend on the average climate of the native habitat. Temperate species (e.g., *Q. robur*) share characters with species that inhabit Mediterranean climates (e.g., *Q. faginea* and *Q. cerris*; Gil-Pelegrín *et al*., 2017).

Evergreen oaks also share common features irrespective of their phylogenetic lineage. These species share harder and thicker leaves, wider bundle sheath extensions, a higher coverage of wider bundle sheath extensions, higher cellulose content and an epidermis with thick cuticles and cell walls that are often lignified. Regarding this second group, some interesting insights emerge. Species of sect. *Ilex* show similar foliar features, regardless of their climate of origin. They occupy quite contrasting climates that range in arid intensity from 0 (e.g., *Q. phyllireoides*) to 261 (e.g., *Q. calliprinus*) (Fig. **5b**) and there is still a great variation when we consider the phylogeny (Fig. **S5**). These shared features may be explained by their common phylogenetic origin in the palaeotropical flora in East Asia (Jiang *et al*., 2019) (Fig. **5a**). Species from this section are currently found in diverse climates, from humid temperate or subtropical climates (e.g., *Q, phillyreoides* and *Q. semecarpifolia*; cf. Jiang *et al*., 2019), to semi-arid Mediterranean areas (e.g., *Q. coccifera*; cf. Vilagrosa *et al*. 2003; Gil-Plegrín *et al*., 2017). According to fossil record, xeromorphic-like traits were already present in the ancestor of the sect. *Ilex*, *Q. yangyiensis* that lived in warm aseasonal tropical conditions (Zhou *et al*., 2007; He *et al*., 2014). Under conditions favorable year-round growth, maintaining an evergreen leaf habit would have been favored over a deciduous leaf habit (Kikuzawa 1991, 1995). Evergreen species with longer leaf lifespan necessitated more durable leaves with traits that allow them to resist tearing and wear due to abiotic (Nikklas, 1999) and biotic (Wrigth & Vincent, 1996; Peeters *et al*., 2007) interactions with the environment (Turner, 1994). The increase in the level of sclerophyll through the accumulation of structural carbohydrates could improve leaf persistence. In this way, the improved leaf protection would help to resist for a longer time (Turner, 1994; Takashima *et al*., 2004). Moreover, habitats with several ecological limitations (e.g., drought, nutrient scarcity, low temperatures during the vegetative period or mechanical damage) are the ones where sclerophylly is more prevalent, as exemplified by the case of Mediterranean evergreens (Alonso-Forn *et al.*, 2020 and references therein).

Although the leaf features of evergreen sclerophylls are extremely functional in Mediterranean-type climate conditions, Ackerly (2004) proposed that these qualities originated in ancestral non-Mediterranean-type habitats in many lineages. Thus, previously mentioned sclerophyllous features can be inferred to have contributed to the later success of section *Ilex* with the expansion of the relatively recent Mediterranean-type climates (Ackerly, 2004; Deng *et al*., 2017). Sect. *Ilex* species may thus have evolved foliar traits under ancestral climates that pre-adapted them to Mediterranean-type climates (Verdú *et al.*, 2003). All American oak clades share a common high latitude temperate ancestor (Hipp *et al*., 2018). Thus, some oaks studied here that occupy habitats in Mediterranean climates in the Nearctic (e.g., *Q. agrifolia* or *Q. wislizeni,* section *Lobatae*), have been suggested to have evolved from temperate ancestors (Hipp *et al.*, 2018). Nevertheless, species of section *Protobalanus* are evergreen and are restricted to mild climates. It appears likely that both conservatism in foliar traits linked to shared ancestry and evolutionary legacies of adaptation to ancient climates as well as evolutionary convergence in foliar traits from different sections contributes to the observed patterns of sclerophyllous traits among lineages and leaf habits. To conclude, there is an evolutionary progression towards sclerophyllous leaf traits in sect. *Protobalanus* and sect. *Lobatae* that inhabit dry climates, while sect. *Ilex* may have evolved to a lesser degree because its tropical ancestors had foliar traits that were pre-adaptated to some extent.

## Acknowledgements

Financial support from Instituto Nacional de Investigación y Tecnología Agraria y Alimentaria (INIA) grant number RTA2015-00054-C02-01, grant PID2019-106701RR-I00 funded by MCIN/AEI/ 10.13039/501100011033 and from Gobierno de Aragón H09_20R research group. Work of DA-F is supported by an FPI-INIA contract BES-2017-081208. Work of MDF is supported by a Juan de la Cierva-Incorporación contract (JC2020-043487-I) provided by MICINN (Spain) and by EU (NextGenerationEU). work of R.M-S. is supported by a predoctoral Gobierno de Aragón scholarship. Work of MN is supported by postdoctoral fellowship Juan de la Cierva-Formación (FJC2020-043902-I), financed by MCIN/AEI/10.13039/501100011033 (Spain) and the European Union (NextGenerationEU/PRTR).

## Author Contribution

DA-F, DS-K, JJP-P, TGA-A and EG-P planned and designed the research. DA-F, DS-K, MDF, RM-S, JJP-P, JPF and EG-P performed the measurements. DA-F, DS-K, MN, JPF and VRD analysed data. DA-F, DS-K, YO and JC-B drafted the manuscript. All authors edited the manuscript with valuable inputs.

## Data availability

Data that support the findings of this study are openly available in TRY at http://doi.org/10.17871/TRY.86.

## Supporting information

**Table S1.**
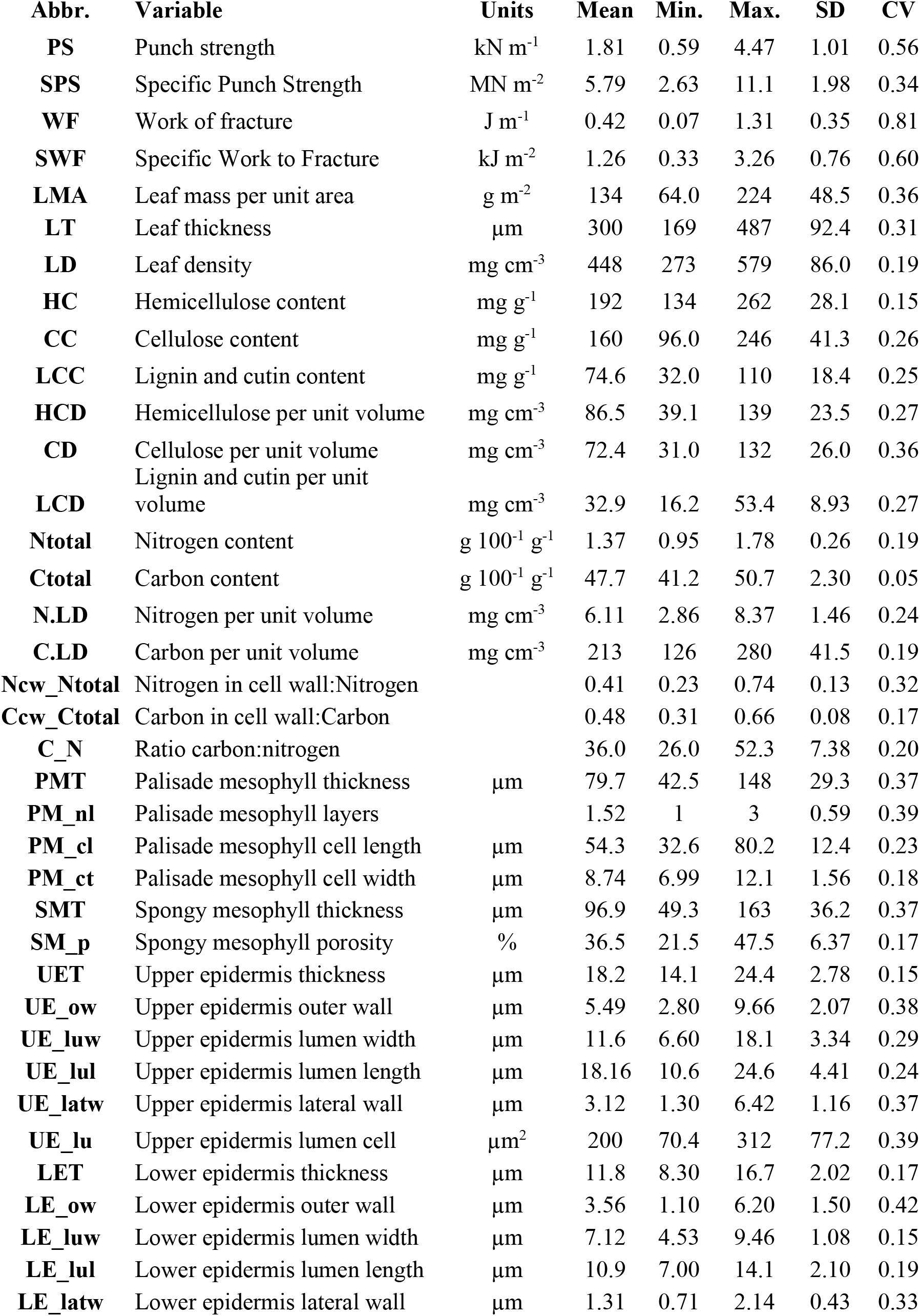
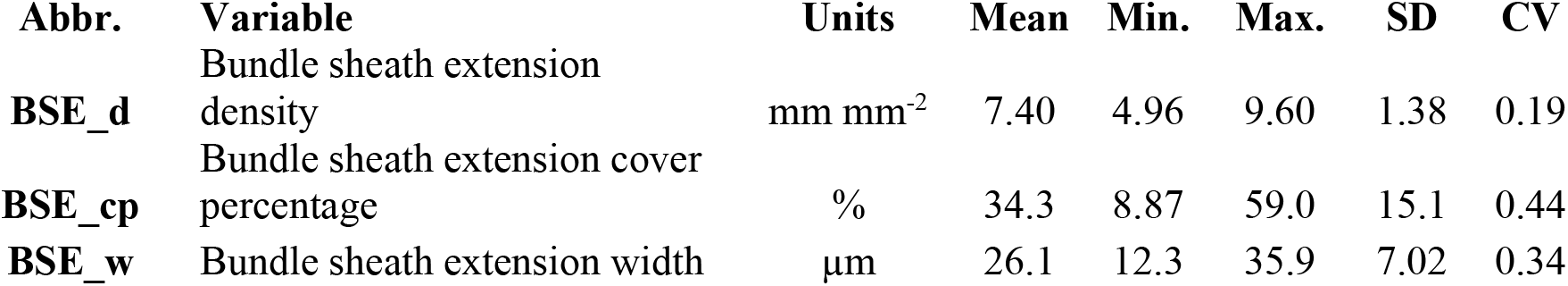
Mean value, maximum (Max.) and minimum (Min.) values, standard deviation (SD) and coefficient of variation (CV) of leaf traits for *Quercus* species.

| Abbr. | Variable | Units | Mean | Min. | Max. | SD | CV |
| --- | --- | --- | --- | --- | --- | --- | --- |
| PS | Punch strength | kN m<sup>-1</sup> | 1.81 | 0.59 | 4.47 | 1.01 | 0.56 |
| SPS | Specific Punch Strength | MN m<sup>-2</sup> | 5.79 | 2.63 | 11.1 | 1.98 | 0.34 |
| WF | Work of fracture | J m<sup>-1</sup> | 0.42 | 0.07 | 1.31 | 0.35 | 0.81 |
| SWF | Specific Work to Fracture | kJ m<sup>-2</sup> | 1.26 | 0.33 | 3.26 | 0.76 | 0.60 |
| LMA | Leaf mass per unit area | g m<sup>-2</sup> | 134 | 64.0 | 224 | 48.5 | 0.36 |
| LT | Leaf thickness | μm | 300 | 169 | 487 | 92.4 | 0.31 |
| LD | Leaf density | mg cm<sup>-3</sup> | 448 | 273 | 579 | 86.0 | 0.19 |
| HC | Hemicellulose content | mg g<sup>-1</sup> | 192 | 134 | 262 | 28.1 | 0.15 |
| CC | Cellulose content | mg g<sup>-1</sup> | 160 | 96.0 | 246 | 41.3 | 0.26 |
| LCC | Lignin and cutin content | mg g<sup>-1</sup> | 74.6 | 32.0 | 110 | 18.4 | 0.25 |
| HCD | Hemicellulose per unit volume | mg cm<sup>-3</sup> | 86.5 | 39.1 | 139 | 23.5 | 0.27 |
| CD | Cellulose per unit volume | mg cm<sup>-3</sup> | 72.4 | 31.0 | 132 | 26.0 | 0.36 |
| LCD | Lignin and cutin per unit volume | mg cm<sup>-3</sup> | 32.9 | 16.2 | 53.4 | 8.93 | 0.27 |
| Ntotal | Nitrogen content | g 100<sup>-1</sup> g<sup>-1</sup> | 1.37 | 0.95 | 1.78 | 0.26 | 0.19 |
| Ctotal | Carbon content | g 100<sup>-1</sup> g<sup>-1</sup> | 47.7 | 41.2 | 50.7 | 2.30 | 0.05 |
| N.LD | Nitrogen per unit volume | mg cm<sup>-3</sup> | 6.11 | 2.86 | 8.37 | 1.46 | 0.24 |
| C.LD | Carbon per unit volume | mg cm<sup>-3</sup> | 213 | 126 | 280 | 41.5 | 0.19 |
| Ncw_Ntotal | Nitrogen in cell wall:Nitrogen |  | 0.41 | 0.23 | 0.74 | 0.13 | 0.32 |
| Ccw_Ctotal | Carbon in cell wall:Carbon |  | 0.48 | 0.31 | 0.66 | 0.08 | 0.17 |
| C_N | Ratio carbon:nitrogen |  | 36.0 | 26.0 | 52.3 | 7.38 | 0.20 |
| PMT | Palisade mesophyll thickness | μm | 79.7 | 42.5 | 148 | 29.3 | 0.37 |
| PM_nl | Palisade mesophyll layers |  | 1.52 | 1 | 3 | 0.59 | 0.39 |
| PM_cl | Palisade mesophyll cell length | μm | 54.3 | 32.6 | 80.2 | 12.4 | 0.23 |
| PM_ct | Palisade mesophyll cell width | μm | 8.74 | 6.99 | 12.1 | 1.56 | 0.18 |
| SMT | Spongy mesophyll thickness | μm | 96.9 | 49.3 | 163 | 36.2 | 0.37 |
| SM_p | Spongy mesophyll porosity | % | 36.5 | 21.5 | 47.5 | 6.37 | 0.17 |
| UET | Upper epidermis thickness | μm | 18.2 | 14.1 | 24.4 | 2.78 | 0.15 |
| UE_ow | Upper epidermis outer wall | μm | 5.49 | 2.80 | 9.66 | 2.07 | 0.38 |
| UE_luw | Upper epidermis lumen width | μm | 11.6 | 6.60 | 18.1 | 3.34 | 0.29 |
| UE_lul | Upper epidermis lumen length | μm | 18.16 | 10.6 | 24.6 | 4.41 | 0.24 |
| UE_latw | Upper epidermis lateral wall | μm | 3.12 | 1.30 | 6.42 | 1.16 | 0.37 |
| UE_lu | Upper epidermis lumen cell | μm<sup>2</sup> | 200 | 70.4 | 312 | 77.2 | 0.39 |
| LET | Lower epidermis thickness | μm | 11.8 | 8.30 | 16.7 | 2.02 | 0.17 |
| LE_ow | Lower epidermis outer wall | μm | 3.56 | 1.10 | 6.20 | 1.50 | 0.42 |
| LE_luw | Lower epidermis lumen width | μm | 7.12 | 4.53 | 9.46 | 1.08 | 0.15 |
| LE_lul | Lower epidermis lumen length | μm | 10.9 | 7.00 | 14.1 | 2.10 | 0.19 |
| LE_latw | Lower epidermis lateral wall | μm | 1.31 | 0.71 | 2.14 | 0.43 | 0.33 |

| **Abbr.** | **Variable** | **Units** | **Mean** | **Min.** | **Max.** | **SD** | **CV** |
| --- | --- | --- | --- | --- | --- | --- | --- |
| **BSE_d** | Bundle sheath extension density | mm mm<sup>-2</sup> | 7.40 | 4.96 | 9.60 | 1.38 | 0.19 |
| **BSE_cp** | Bundle sheath extension cover percentage | % | 34.3 | 8.87 | 59.0 | 15.1 | 0.44 |
| **BSE_w** | Bundle sheath extension width | μm | 26.1 | 12.3 | 35.9 | 7.02 | 0.34 |

**Table S2.** Heat of combustion, ash, nitrogen concentration (N) and construction cost (CC) of leaves for the seven studied *Quercus* species. Data are mean ± SE. Different letters indicate significant differences among species (Tukey’s test, *P* < 0.05).

| Species | Heat of combustion<br>(kJ g<sup>-1</sup>) | Ash<br>(g g<sup>-1</sup>) | N<br>(g g<sup>-1</sup>) | CC<br>(g glucose g<sup>-1</sup>) |
| --- | --- | --- | --- | --- |
| *Q. chrysolepis* | 22.2 ± 0.3<sup>b</sup> | 0.0736 ± 0.0004<sup>c</sup> | 0.012 ± 0.001<sup>a</sup> | 1.58 ± 0.02<sup>bc</sup> |
| *Q. agrifolia* | 21.9 ± 0.3<sup>ab</sup> | 0.0465 ± 0.0004<sup>b</sup> | 0.014 ± 0.002<sup>a</sup> | 1.61 ± 0.02<sup>bc</sup> |
| *Q. wislizeni* | 19.8 ± 0.6<sup>a</sup> | 0.0392 ± 0.0005<sup>b</sup> | 0.014 ± 0.003<sup>a</sup> | 1.46 ± 0.04<sup>a</sup> |
| *Q. coccifera* | 22.0 ± 0.9<sup>ab</sup> | 0.0487 ± 0.0002<sup>b</sup> | 0.014 ± 0.002<sup>a</sup> | 1.61 ± 0.06<sup>bc</sup> |
| *Q. ilex* subsp. *ilex* | 20.8 ± 2.1<sup>ab</sup> | 0.0558 ± 0.0006<sup>bc</sup> | 0.011 ± 0.003<sup>a</sup> | 1.50 ± 0.16<sup>abc</sup> |
| *Q. ilex* subsp. *rotundifolia* | 20.0 ± 0.7<sup>a</sup> | 0.0257 ± 0.0004<sup>a</sup> | 0.011 ± 0.001<sup>a</sup> | 1.49 ± 0.06<sup>ab</sup> |
| *Q. suber* | 20.3 ± 0.7<sup>a</sup> | 0.0856 ± 0.0004<sup>c</sup> | 0.017 ± 0.002<sup>a</sup> | 1.43 ± 0.05<sup>a</sup> |

**Fig. S1.**
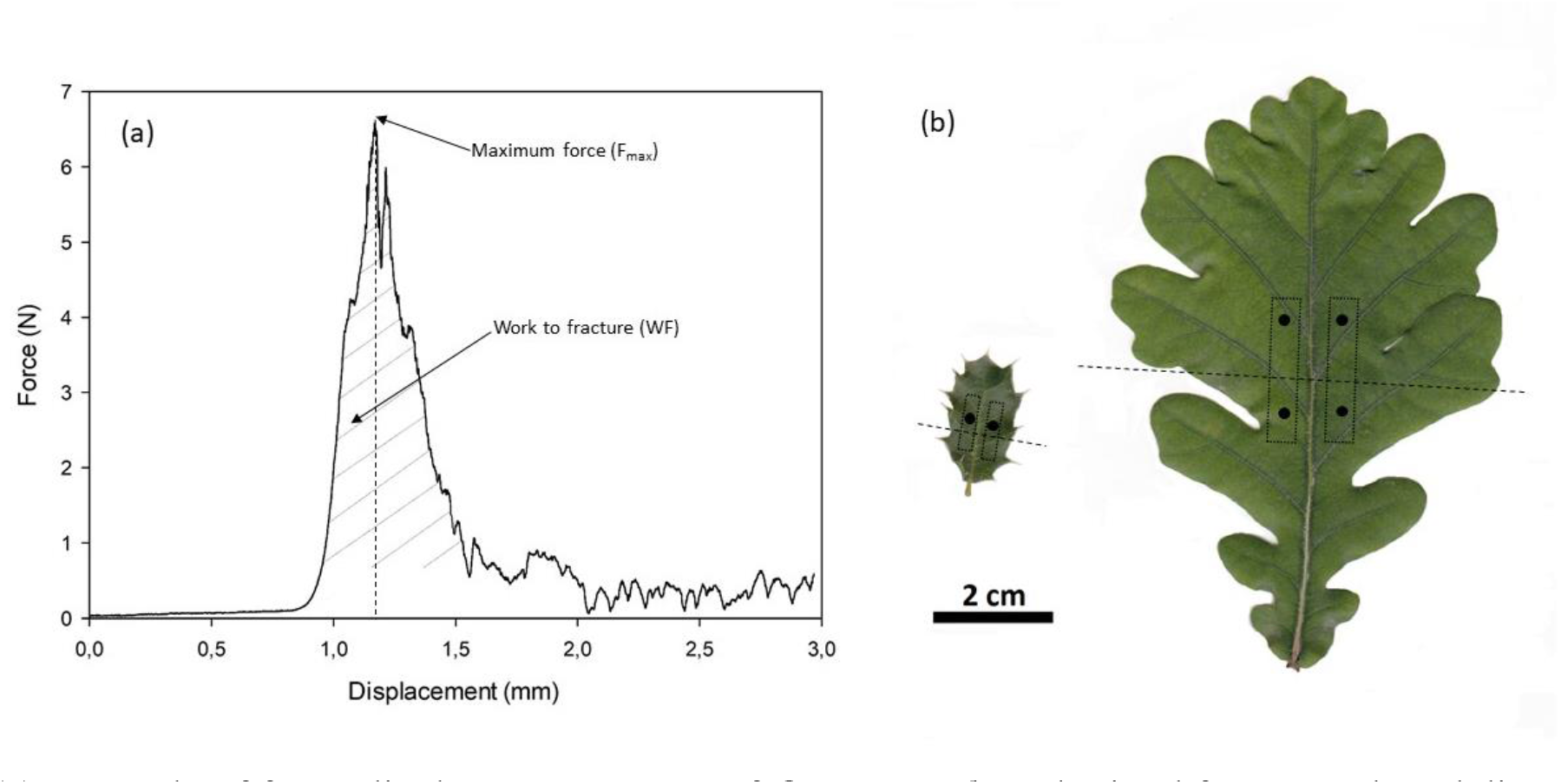
(a) Example of force-displacement curves of *Quercus robur* obtained from punch and die test showing maximum force (F_max_) and work to fracture (WF) calculated as the area below the curve between the initial contact of the punch with the leaf and F_max_. (b) Examples of the points were punch and die tests were conducted in leaves of *Q. coccifera* (left) and *Q. robur* (right), avoiding major veins and delimiting upper and lower borders of the intercostal panel.

**Fig. S2.**
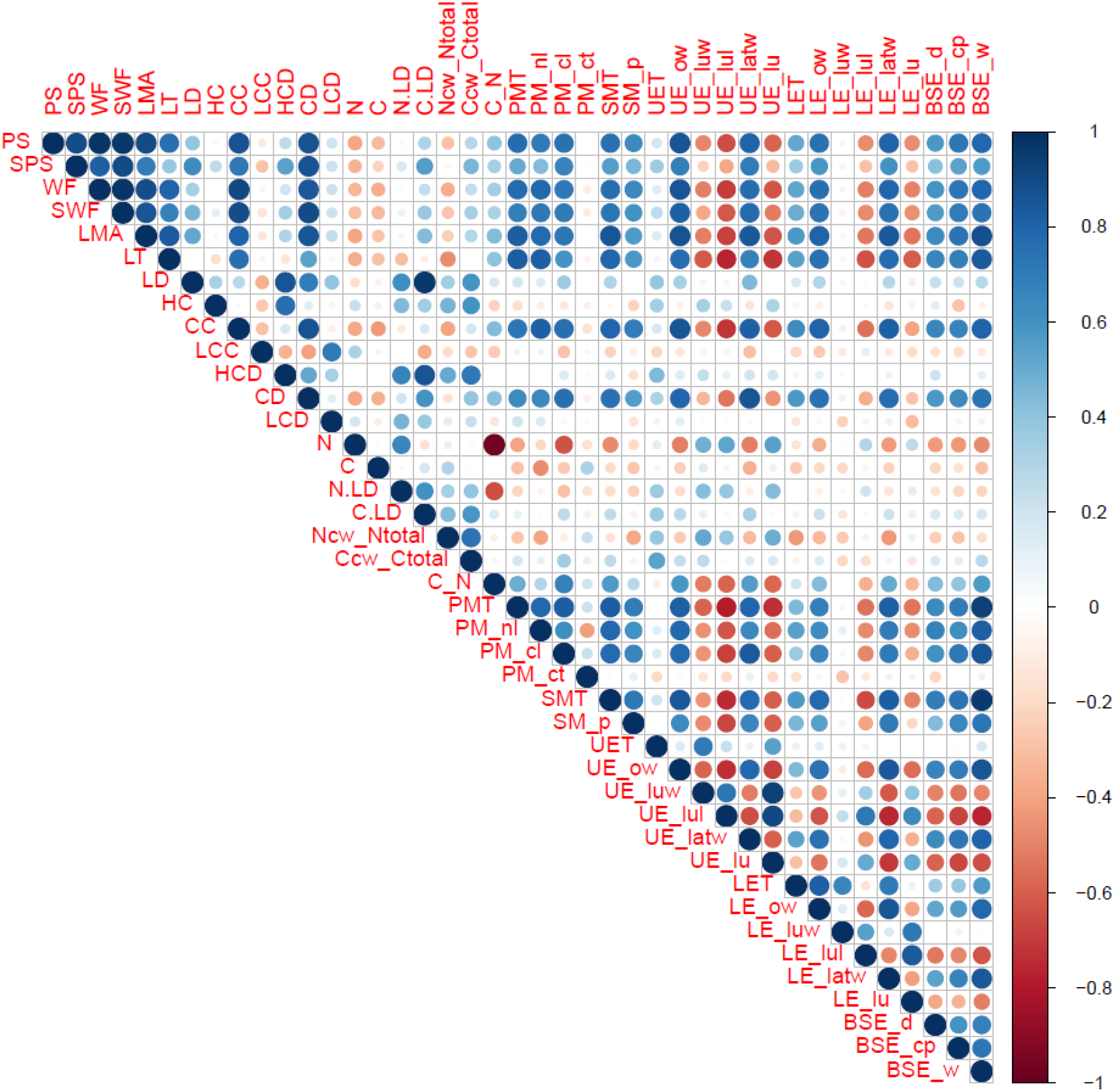
Correlation coefficients in matrix form between leaf mechanical properties and anatomical traits. The matrix includes the correlations graduated according to the legend.

**Fig. S3.**
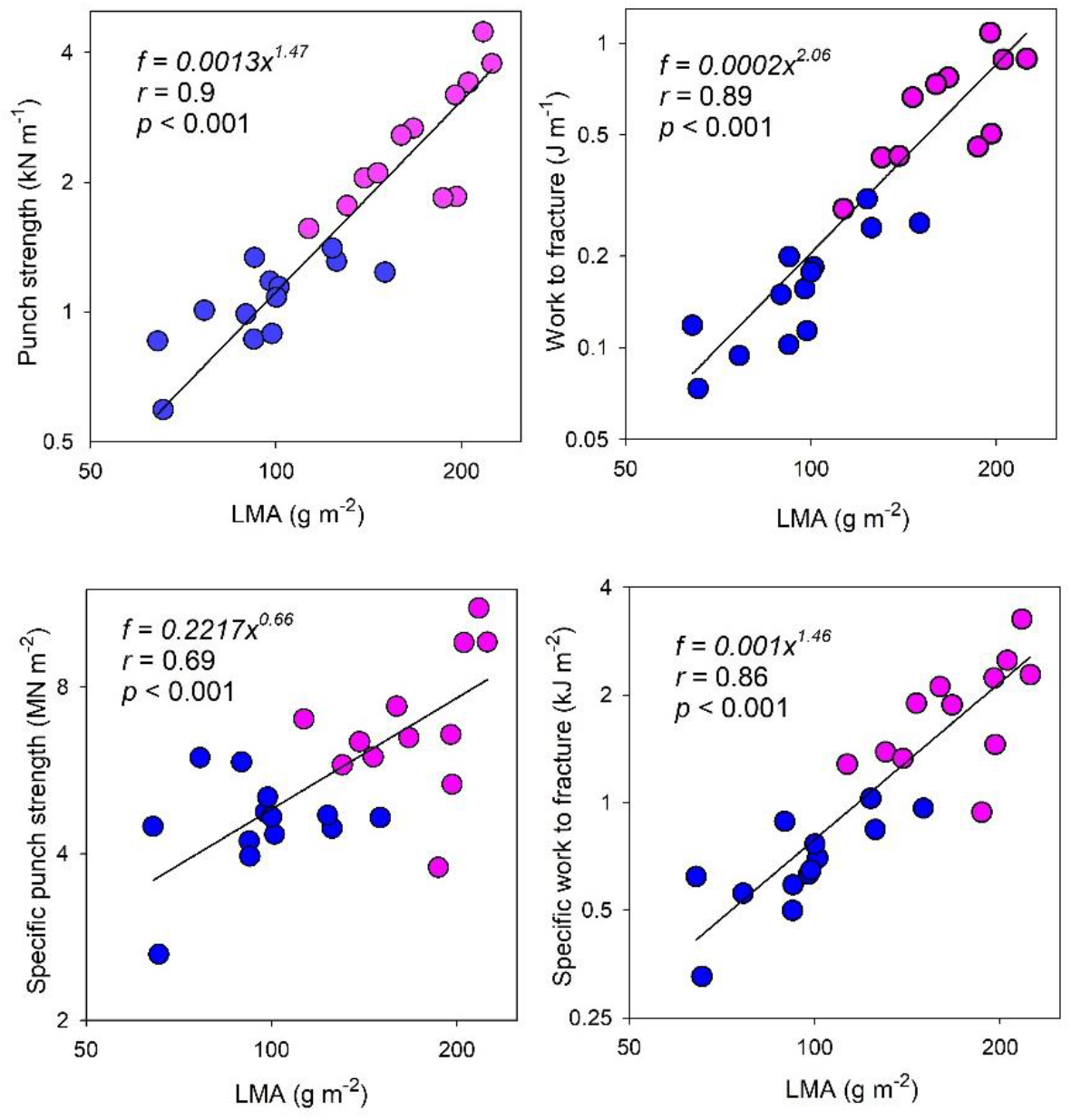
The bivariate trait relationships between leaf structural properties and leaf mass per area (LMA) for deciduous (DEC; blue) and evergreen (EVE; pink) *Quercus* species. Note that graph axes are log_10_ scaled.

**Fig. S4.**
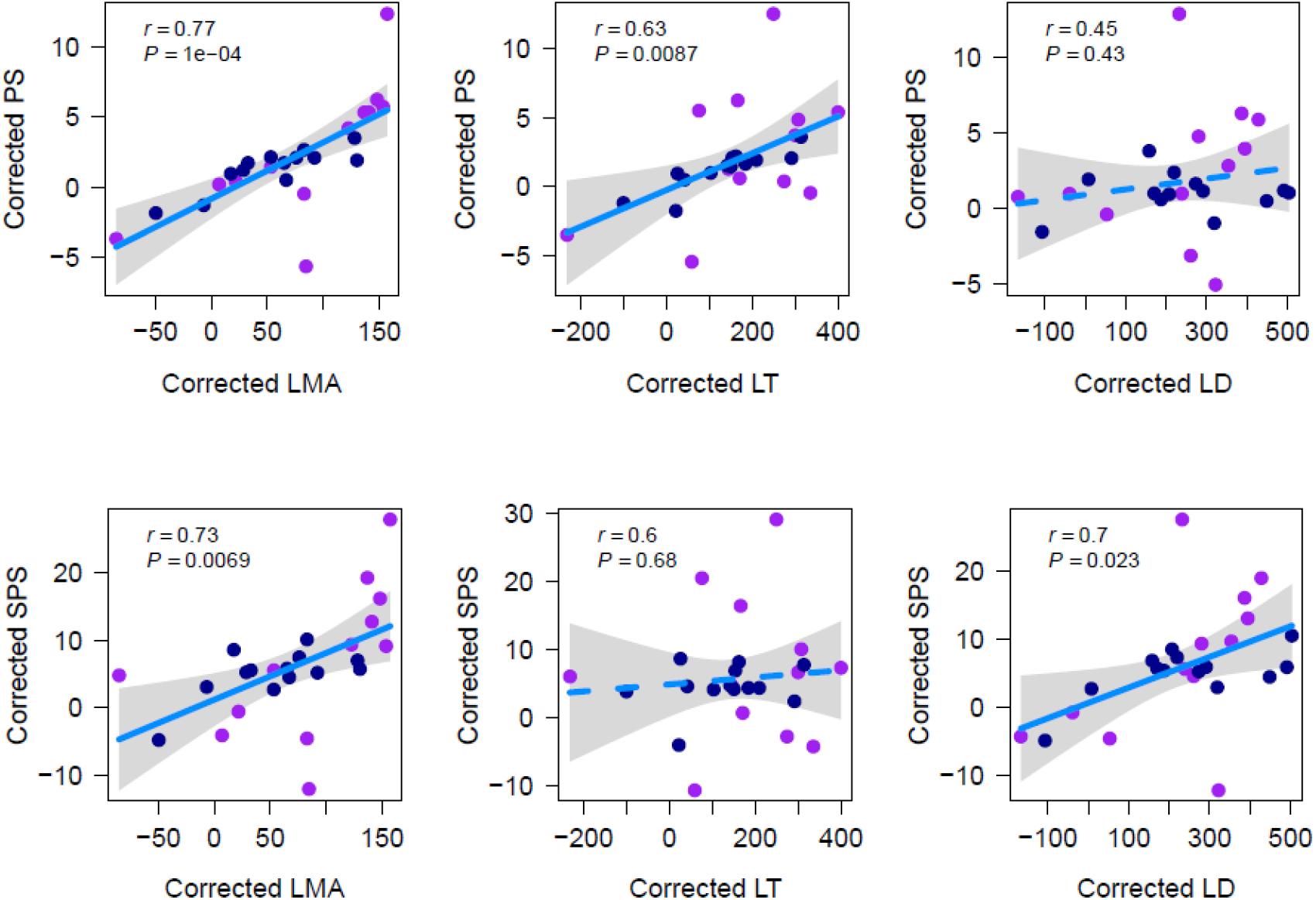
Corrected main relationships using phylogenetic generalized least squares analysis assuming that trait evolution mimics Brownian motion and using the phylogeny from Hipp *et al*. (2020). Physical parameters (punch strength and specific punch strength) and leaf mass per area, leaf thickness, and leaf density, for deciduous (DEC; blue) and evergreen (EVE; fuchsia) *Quercus* species. Each circle belongs to a Quercus species and represents its mean value. The blue continuous line is the correlation considering all species.

**Fig. S5.**
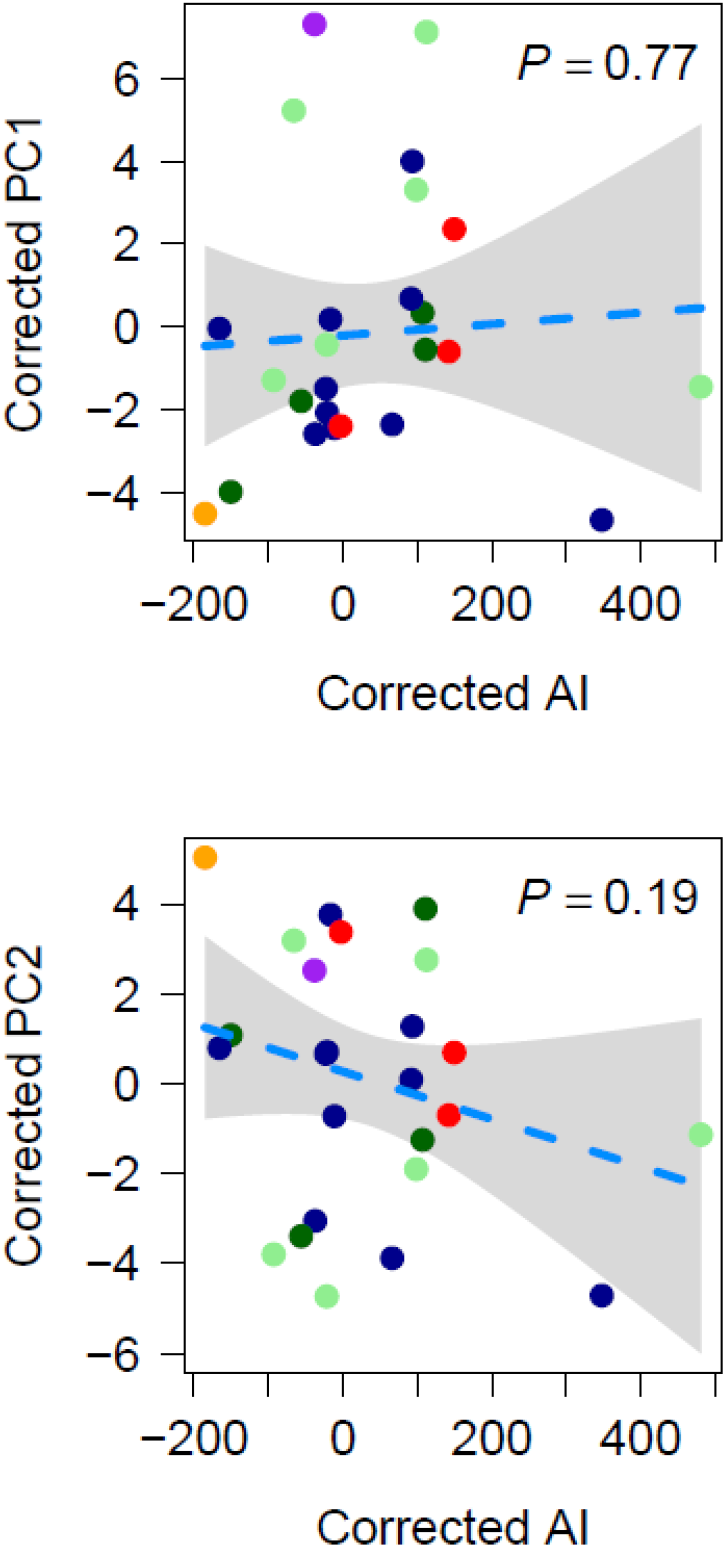
Corrected relationship between PC1 and PC2 and Arid intensity (AI) using phylogenetic generalized least squares analysis assuming that trait evolution mimics Brownian motion and using the phylogeny from Hipp *et al.* (2020). Each circle belongs to a *Quercus* species and represents its mean value. The colors of the circles represent the different sections of studied *Quercus* (see Figure 5).

## References

Alonso-Forn D, Sancho-Knapik D, Ferrio JP, Peguero-Pina JJ, Bueno A, Onoda Y, Cavender-Bares J, Ülo N, Jansen S, Riederer M et al. 2020. Revisiting the functional basis of sclerophylly within the leaf economics spectrum of oaks: different roads to Rome. Current Forestry Reports 6: 260–281.

Ackerly D. 2004. Functional strategies of chaparral shrubs in relation to seasonal water deficit and disturbance. Ecological Monographs 74: 25–44.

Aranwela N, Sanson G, Read J. 1999. Methods of assessing leaf-fracture properties. New Phytologist 144: 369–393.

Beadle NCW. 1966. Soil phosphate and its role in molding segments of the australian flora and vegetation, with special reference to xeromorphy and sclerophylly. Ecology 47: 992–1007.

Bennett RN, Wallsgrove RM. 1994. Secondary metabolites in plants defense mechanisms. New Phytologist 127: 617–633.

Carver TLW, Gurr SJ. 2006. Filamentous fungi on plant surfaces. Pages 368 397 in: Biology of the Plant Cuticle. Riederer M and Müller M, eds. Blackwell Publishing Ltd., Oxford.

Chabot BF, Hicks DJ. 1982. The ecology of leaf life spans. Annual Review of Ecology and Systematics 13: 229–259.

Cherrett JM. 1968. A Simple Penetrometer for Measuring Leaf Toughness in Insect Feeding Studies. Journal of Economic Entomology 61: 1736–1738.

Choong MF, Lucas PW, Ong JSY, Pereira B, Tan HTW, Turner IM. 1992. Leaf fracture toughness and sclerophylly: their correlations and ecological implications. New Phytologist 121: 597–610.

Coley PD. 1983. Herbivory and Defensive Characteristics of Tree Species in a Lowland Tropical Forest. Ecology Monographs 53: 209–234.

Coley PD, Barone JA. 1996. Herbivory and plant defenses in tropical forests. Annual Review of Ecology and Systematics 27: 305–335.

Edwards C, Read J, Sanson G. 2000. Characterising sclerophylly: Some mechanical properties of leaves from heath and forest. Oecologia 123: 158–167.

Feeny P. 1970. Seasonal Changes in Oak Leaf Tannins and Nutrients as a Cause of Spring Feeding by Winter Moth Caterpillars. Ecology 51: 565–581.

Gil-Pelegrín E, Saz MA, Cuadrat JM, Peguero-Pina JJ, Sancho-Knapik D. 2017. Oaks under Mediterranean-type climates: functional response to summer aridity. In: Gil-Pelegrín E, Peguero-Pina JJ, Sancho-Knapik D, eds. Oaks Physiological Ecology. Exploring the Functional Diversity of Genus Quercus L. Springer International Publishing AG, Cham, Switzerland.

Goering HK, Van Soest PJ. 1970. Forage fiber analyses (apparatus, reagents, procedures, and some applications). USDA–ARS Agriculture Handbook, U.S. Gov. Print. Office, Washington, DC.

Gonçalves-Alvim SJ, Korndorf G, Fernandes GW. 2006. Sclerophylly in *Qualea parviflora* (Vochysiaceae): Influence of herbivory, mineral nutrients, and water status. Plant Ecology 187: 153–162.

Grubb PJ. 1986. Sclerophylls, pachyphylls and pycnophylls: the nature and significance of hard leaf surfaces. In: Juniper B, Southwood R, eds. Insects and the plant surface. London: Edward Arnold.

Harborne JB. 1990. Constraints on the evolution of biochemical pathways. Journal of Linnean Society 39: 135–151.

Hennig C, Liao TF. 2013. How to find an appropriate clustering for mixed-type variables with application to socio-economic stratification. Journal of the Royal Statistical Society: Series C (Applied Statistics) 62: 309–369.

Hipp AL, Manos PS, Hahn M, Avishai M, Bodénès C, Cavender-Bares J, Crowl AA, Deng M, Denk T, Fitz-Gibbon S et al. 2020. Genomic landscape of the global oak phylogeny. New Phytologist 226: 1198–1212.

Jiang X-L, Hipp AL, Deng M, Su T, Zhou Z-K, Yan M-X. 2019. East Asian origins of European holly oaks (Quercus section Ilex Loudon) via the Tibet-Himalaya. Journal of Biogeography 46: 2203–2214.

Johansen, D.A. 1940. Plant Microtechnique. McGraw-Hill, New York.

John GP, Scoffoni C, Buckley TN, Villar R, Poorter H, Sack L. 2017. The anatomical and compositional basis of leaf mass per area. Ecology Letters 20: 412–425.

Lê S, Josse J, Husson F. 2008. FactoMineR: an R package for multivariate analysis. Journal of Statistical Software 25: 1–18.

Kaufman L, Rousseeuw PJ. 1987. Clustering by means of medoids. In: Dodge Y, ed. Statistical Data Analysis Based on the L1 Norm and Related Methods, Basel: Birkhäuser Publishing.

Kikuzawa, K. 1991. A cost-benefit analysis of leaf habit and leaf longevity of trees and their geographical patterns. The American Naturalist 138: 1250–1263.

Kikuzawa, K. 1995. The basis for variation in leaf longevity of plants. Vegetatio 121: 89–100.

Kikuzawa K, Onoda Y, Wright IJ, Reich PB. 2013. Mechanisms underlying global temperature-related patterns in leaf longevity. Global Ecology and Biogeography 22: 982–993.

Kitajima K, Poorter L. 2010. Tissue-level leaf toughness, but not lamina thickness, predicts sapling leaf lifespan and shade tolerance of tropical tree species. New Phytologist 186: 708–721.

Koppel A, Heinsoo K. 1994. Variability in cuticular resistance of *Picea abies* (L.) karst. and its significance in winter desiccation. Proceedings of the Estonian Academy of Sciences 4: 56–63.

Kouki M, Manetas Y. 2002. Toughness is less important than chemical composition of Arbutus leaves in food selection by *Poecilimon* species. New Phytologist 154: 399–407.

Krauss P, Markstädter C, Riederer M. 1997. Attenuation of UV radiation by plant cuticles from woody species. Plant, Cell and Environment 20: 1079–1085.

Lamontagne M, Margolis H, Bigras F. 1998. Photosynthesis of black spruce, jack pine, and trembling aspen after artificially induced frost during the growing season. Canadian Journal of Forest Research 28: 1–12.

Loveless AR. 1961. A nutritional interpretation of sclerophylly based on differences in the chemical composition of sclerophyllous and mesophytic leaves. Annals of Botany 25: 168–184.

Loveless AR. 1962. Further evidence to support a nutritional interpretation of Sclerophylly. Annals of Botany 26: 551–560.

Matsuki S, Koike T. 2006. Comparison of Leaf Life Span, Photosynthesis and Defensive Traits Across Seven Species of Deciduous Broad-leaf Tree Seedlings. Annals of Botany 97: 813–817.

Makkar HPS, Bluemmel M, Borowy NK, Becker K. 1993. Gravimetric determination of tannins and their correlations with chemical and protein precipitation methods. Journal of the Science of Food and Agriculture 61: 161–165.

Mediavilla S, Garcia-Ciudad A, Garcia-Criado B, Escudero A. 2008. Testing the correlations between leaf life span and leaf structural reinforcement in 13 species of European Mediterranean woody plants. Functional Ecology 22: 787–793.

Mediavilla S, Babiano J, Martínez-Ortega M, Escudero A. 2018. Ontogenetic changes in anti-herbivore defensive traits in leaves of four Mediterranean co-occurring *Quercus* species. Ecological Research 33: 1093–1102.

Moles AT, Peco B, Wallis IR, Foley WJ, Poore AGB, Seabloome EW, Vesk PA, Bisigato AJ, Cella-Pizarro L, Clark CJ et al. 2013. Correlations between physical and chemical defences in plants: tradeoffs, syndromes, or just many different ways to skin a herbivorous cat? New Phytologist 198: 252–263.

Niinemets Ü. 1997. Energy requirement for foliage construction depends on tree size in young *Picea abies* trees. Trees – Structure and Function 11: 420–431.

Niinemets Ü. 1999. Research review. Components of leaf dry mass per area - thickness and density – alter leaf photosynthetic capacity in reverse directions in woody plants. New Phytologist 144: 35–47.

Niinemets Ü. 2001. Global-scale climatic controls of leaf dry mass per area, density, and thickness in trees and shrubs. Ecology 82: 453–469.

Niklas KJ. 1999. A mechanical perspective on foliage leaf form and function. New Phytologist 143: 19–31.

Nikolopoulos D, Liakopoulos G, Drossopoulos I, Karabourniotis G. 2002. The relationship between anatomy and photosynthetic performance of heterobaric leaves. Plant Physiologist 129: 235–243.

Onoda Y, Westoby M, Adler PB, Choong AMF, Clissold FJ, Cornelissen JHC, Díaz S, Dominy NJ, Elgart A, Enrico L et al. 2011. Global patterns of leaf mechanical properties. Ecology Letters 14: 301–312.

Onoda Y, Richards L, Westoby M. 2012. The importance of leaf cuticle for carbon economy and mechanical strength. New Phytologist 196: 441–447.

Onoda Y, Schieving F, Anten NPR. 2015. A novel method of measuring leaf epidermis and mesophyll stiffness shows the ubiquitous nature of the sandwich structure of leaf laminas in broad-leaved angiosperm species. Journal of Experimental Botany 66: 2487–2499.

Oertli JJ, Lips SH, Agami M. 1990. The strength of sclerophyllous cells to resist collapse due to negative turgot pressure. Acta Oecologica 11: 281–290.

Peeters PJ. 2002. Correlations between leaf structural traits and the densities of herbivorous insect guilds. Biological Journal of the Linnean Society 77: 43–65.

Peeters PJ, Sanson G, Read J. 2007. Leaf biomechanical properties and the densities of herbivorous insect guilds. Functional Ecology 21: 246–255.

Peguero-Pina JJ, Aranda I, Cano FJ, Galmés J, Gil-Pelegrín E, Niinemets U, Sancho-Knapik D, Flexas J. 2017. The Role of Mesophyll Conductance in Oak Photosynthesis: Among- and Within-Species Variability. In: Oaks Physiological Ecology. Exploring the Functional Diversity of Genus Quercus L. Gil-Pelegrín E, Peguero-Pina JJ, Sancho-Knapik D, eds. Springer International Publishing.

Pérez-Harguindeguy N, Díaz S, Vendramini F, Cornelissen JHC, Gurvich DE, Cabido M. 2003. Leaf traits and herbivore selection in the field and in cafeteria experiments. Austral Ecology 28: 642–650.

Phillipson J. 1964. A miniature bomb calorimeter for small biological samples. Oikos 15: 130–139.

Porter LJ, Hrstich LN, Chan BG. 1986. The conversion of procyanidins and prodelphinidins to cyanidin and delphinidin. Phytochemistry 25: 223–230.

Poorter H, Niinemets Ü, Poorter L, Wright IJ, Villar R. 2009. Causes and consequences of variation in leaf mass per area (LMA): A meta-analysis. New Phytologist 182: 565–588.

Poorter H, Villar R. 1997. The fate of acquired carbon in plants: chemical composition and construction costs. In: Bazzaz FA, Grace J, eds. Plant resource allocation. San Diego, CA, USA: Academic Press.

Pradhan Mitra P, Loqué D. 2014. Histochemical staining of Arabidopsis thaliana secondary cell wall elements. Journal of Visualized Experiments 87: 51381.

R Core Team. 2021. R: A language and environment for statistical computing. R Foundation for Statistical Computing, Vienna, Austria. URL https://www.R-project.org/.

Rodríguez-Calcerrada J, Sancho-Knapik D, Martin-StPaul NK, Limousin JM, McDowell NG, Gil-Pelegrín E. 2017. Drought-Induced Oak Decline—Factors Involved, Physiological Dysfunctions, and Potential Attenuation by Forestry Practices. In: Oaks Physiological Ecology. Exploring the Functional Diversity of Genus Quercus L. Gil-Pelegrín E, Peguero-Pina JJ, Sancho-Knapik D, eds. Springer International Publishing.

Rosseel, Y. 2012. lavaan: An R Package for Structural Equation Modeling. Journal of Statistical Software 48: 1–36.

Sancho-Knapik D, Escudero A, Mediavilla S, Scoffoni C, Zailaa J, Cavender-Bares J, Álvarez-Arenas TG, Molins A, Alonso-Forn D, Ferrio JP et al. 2021. Deciduous and evergreen oaks show contrasting adaptive responses in leaf mass per area across environments. New Phytologist 230: 521–534.

Scoffoni C, Rawls M, McKown A, Cochard H, Sack L. 2011. Decline of leaf hydraulic conductance with dehydration: relationship to leaf size and venation architecture. Plant Physiology 156: 832–843.

Thomas FM, Blank R, Hartmann G. 2002. Abiotic and biotic factors and their interactions as causes of oak decline in Central Europe. Forest Pathology 32: 277–307.

Turner IM. 1994. Sclerophylly: primarily protective? Functional Ecology 8: 669–675.

Verdú M, Dávila P, García-Fayos P, Flores-Hernández N, Valiente-Banuet A. 2003. ‘Convergent’ traits of mediterranean woody plants belong to pre-mediterranean lineages, Biological Journal of the Linnean Society 78: 415–427.

Vilagrosa A, Bellot J, Vallejo VR, Gil-Pelegrín E. 2003. Cavitation, stomatal conductance, and leaf dieback in seedlings of two co-occurring Mediterranean shrubs during an intense drought. Journal of Experimental Botany 54: 2015–2024.

Villar R, Merino J. 2001. Comparison of leaf construction costs in woody species with differing leaf life-spans in contrasting ecosystems. New Phytologist 151: 213–226.

Warton DI, Wright IJ, Falster DS, Westoby M. 2006. Bivariate line-fitting methods for allometry. Biol Rev Camb Philos Soc. 81(2):259–91. doi: 10.1017/S1464793106007007.

Wei T, Simko V. 2021. R package ‘corrplot’: Visualization of a Correlation Matrix. (Version 0.92), https://github.com/taiyun/corrplot.

Williams LH. 1954. The Feeding Habits and Food Preferences of Acrididae and the Factors Which Determine Them. Transactions of the Entomological Society of London. 105: 423–454.

Williams K, Percival F, Merino J, Mooney HA. 1987. Estimation of tissue construction cost from heat of combustion and organic nitrogen content. Plant Cell & Environment 10: 725–734.

Wright IJ, Cannon K. 2001. Relationships between leaf lifespan and structural defences in a low-nutrient, sclerophyll flora. Functional Ecology 15: 351–359.

Wright W, Vincent JFV. 1996. Herbivory and the mechanics of fracture in plants. Biological Reviews 71: 401–413.

Wright IJ, Reich PB, Westoby M, Ackerly DD, Baruch Z, Bongers F, Cavender-Bares J, Chapin T, Cornelissen JHC, Diemer M et al. 2004. The worldwide leaf economics spectrum. Nature 428: 821–827.

Zhou Z, Yang Q, Xia K. 2007. Fossils of Quercus sect. Heterobalanus can help explain the uplift of the Himalayas. Chinese Science Bulletin 52: 238–247.

